# The single *Marchantia polymorpha FERONIA* homolog reveals an ancestral role in regulating cellular expansion and integrity

**DOI:** 10.1101/2020.12.23.424085

**Authors:** Martin A. Mecchia, Moritz Rövekamp, Alejandro Giraldo-Fonseca, Dario Meier, Philippe Gadient, Hannes Vogler, Daria Limacher, John L. Bowman, Ueli Grossniklaus

## Abstract

Plant cells are surrounded by a cell wall, a rigid structure that is not only important for cell and organ shape, but crucial for intercellular communication and interactions with the environment. In the flowering plant *Arabidopsis thaliana*, the 17 members of the *Catharanthus roseus* RLK1-like (*Cr*RLK1L) receptor kinase family are involved in a multitude of physiological and developmental processes, making it difficult to assess their primary or ancestral function. To reduce genetic complexity, we characterized the single *Cr*RLK1L gene of *Marchantia polymorpha*, Mp*FERONIA* (Mp*FER*). Plants with reduced Mp*FER* levels show defects in vegetative development, i.e., rhizoid formation and cell expansion, and have reduced male fertility. In contrast, Mp*fer* null mutants and overexpression lines severely affect cell integrity and morphogenesis of the gametophyte. Thus, the *Cr*RLK1L gene family originated from a single gene with an ancestral function in cell expansion and the maintenance of cellular integrity. During land plant evolution, this ancestral gene diversified to fulfil a multitude of specialized physiological and developmental and roles in the formation of both gametophytic and sporophytic structures essential to the life cycle of flowering plants.

**Summary statement:** The *Cr*RLK1L family arose in land plants and the *Marchantia polymorpha genome* contains a single copy, Mp*FER,* which is broadly expressed and regulates cell expansion and cell wall integrity.

## Introduction

The functioning of plant cells is contingent upon the cell wall acting as a barrier between the cell and its environment. This complex matrix of polysaccharides, glycoproteins, and other organic compounds defines the growth and shape of a cell and provides protection against biotic and abiotic stresses [1], [2], [3]. Moreover, the cell wall has to resist turgor pressure while also allowing cell expansion during growth. Thus, sensing and controlling cell wall integrity (CWI) is crucial for plant cells, and receptor kinases (RKs) play an important role in sensing extracellular cues that activate intracellular pathways. Different RK subfamilies are defined by their extracellular domain (ECD), which is more variable than their transmembrane and kinase domains [4]. Over the last decade, the *Cr*RLK1L subfamily emerged as important CWI sensors [5], [6].

Functions have been assigned to 15 of the 17 *Cr*RLK1L members encoded in the *A. thaliana* genome: they are required for sensing environmental cues and cell-cell communication in diverse contexts, including reproduction, hormone signaling, cell expansion, innate immunity, and various stress responses [7]–[23]. For instance, *FERONIA* (At*FER*), *THESEUS1* (At*THE1*), *HERCULES1* (At*HERK1*), and At*HERK2* are required for cell elongation as At*fer* single, At*the1*At*herk1* double, and At*the1*At*herk1*At*herk2* triple mutants exhibit stunted growth [8], [10]–[12]. Root hairs burst in mutants affecting At*FER* and *[Ca^2+^]cyt-ASSOCIATED PROTEIN KINASE1* (At*CAP1*)/*ERULUS* (At*ERU*) [13], [17]. Moreover, At*FER* plays a role in powdery mildew resistance [12], innate immunity [16], [23], calcium signaling [24], phytohormone signaling [11], [25]–[27], and mechano- and heavy metal sensing [21], [28]. In most contexts, the *Cr*RLK1Ls tend to promote cellular growth and cell elongation. In contrast, At*THE1* actively inhibits cell elongation in hypocotyls when cell wall perturbations occur, *e.g.*, due to mutations in the cell wall biosynthesis machinery or upon treatment with isoxaben, a cellulose synthesis inhibitor [8], [29].

In addition to regulating cellular growth during the vegetative phase of the life cycle, several *Cr*RLK1Ls play a role during fertilization by controlling the growth and reception of the pollen tube. In the female gametophyte, At*FER* is highly expressed in the two synergid cells that flank the egg cell and are important for double fertilization [7], [30]. At*FER* is crucial for pollen tube reception and the release of the two sperm cells [7], [11], [24], [31], [32], a process also involving two other, redundantly acting synergid-expressed *Cr*RLK1Ls, *AtHERK1* and *ANJEA* (*AtANJ*) [18]. Moreover, *AtFER* and *AtANJ* also play a role in pollen-stigma recognition during pollen germination [1]. In contrast to these synergid-expressed *Cr*RLK1Ls, four pollen-expressed *Cr*RLK1Ls, named *ANXUR1* (At*ANX1*), At*ANX2, BUDDHA’S PAPER SEAL1* (At*BUPS1*) and At*BUPS2* [9], [15], [19], [33], are redundantly required for pollen tube integrity and tip growth.

To transduce extracellular cues, *Cr*RLK1Ls bind secreted peptides of the RAPID ALKALINIZATION FACTOR (RALF) family [23], [33]–[36], which induce complex formation with members of LORELEI (LRE) family of GPI-anchored proteins [33], [36]– [38]. Hence, it was suggested that at least some *Cr*RLK1Ls act as CWI sensors that coordinate cell elongation in response to changes in the cell wall [39]–[44]. A role as CWI sensors is also consistent with recent findings that AtFER interacts with pectin [43], [45], [46].

The 17 *A. thaliana Cr*RLK1Ls act partially redundantly and sometimes even have opposite effects on cellular growth (reviewed in [20], [42], [44], [47]). To investigate the original function of *Cr*RLK1Ls, we characterized them in a system with reduced genetic complexity: the liverwort *Marchantia polymorpha* encodes a single *Cr*RLK1L [48], [49]. Liverworts represent an early diverging land plant lineage and have been hypothesized to retain, at least in part, characteristics of the earliest land plants [48], [50]–[55]. The presence of similar types of gene families in all plant lineages suggests that the differences in development evolved by co-opting and modifying existing developmental programs and genetic networks, rather than through the evolution of novel genes [50], [51], [56]. *M. polymorpha* is thus an ideal system to study the ancestral role of genes and how it was modulated and diversified during land plant evolution [48], [57], [58].

We refer to the single *Cr*RLK1L encoded in the *M. polymorpha* genome, Mp4g15890 (Mapoly0869s0001), as Mp*FERONIA* (Mp*FER*) [47]–[49] (aka Mp*THESEUS,* Mp*THE*) [59]. An Mp*fer/the* mutant was identified in a large T-DNA screen for defective rhizoid formation, developing short and irregularly shaped rhizoids with brown tips, indicating rhizoid rupture [59]. In the same screen, a mutant affecting the ortholog of the *A. thaliana* receptor-like cytoplasmic kinase *MARIS* (At*MRI*), known to act downstream of At*FER*, was identified with a similar phenotype (named Mp*mri* aka Mp*pti*, Mp7g17560 [Mapoly0051s0094]) [59], [60]. As in *A. thaliana*, a dominant active version of Mp*MRI* can partially rescue the Mp*fer* rhizoid phenotype [60]. This suggests that at least some of the machinery associated with cell elongation and CWI sensing is conserved between *A. thaliana* and *M. polymorpha*.

Here, we report that Mp*FER* function is required for a variety of aspects of *M. polymorpha* gametophyte development. Phylogenetic analyses suggest that *Cr*RLK1L genes first appeared in the common ancestor of land plants. The characterization of lines with reduced Mp*FER* levels indicates that, in addition to its function in rhizoid formation, Mp*FER* controls cell size and organ growth and is involved in male gametogenesis and fertility. Analysis of the Mp*FER* expression pattern points to an involvement in female sexual organogenesis and sporophyte development. Our data suggests that the broad involvement of *Cr*RLK1L function in the development and physiology of land plants is an ancestral and conserved characteristic, although the functions of *Cr*RLK1Ls were extended and adapted to control additional developmental processes in the sporophyte during the course of land plant evolution.

## Results

### *Cr*RLK1L is conserved in land plants and arose in this lineage

The *M. polymorpha* genome encodes a single *Cr*RLK1L homolog (Mp4g15890 [Mapoly0869s0001]) [47]–[49]. Like other *Cr*RLK1L members, this gene encodes an intracellular serine/threonine kinase domain, a transmembrane domain, and an extracellular malectin-like domain with two malectin domains (Fig. 1A). Given that the ECD of *Cr*RLK1Ls in *A. thaliana* is crucial for their function and, in contrast to the kinase domain, not interchangeable [61], we performed a phylogenetic analysis with the amino acid sequences of the predicted malectin-like domain of Mp4g15890, the *Cr*RLK1Ls of the mosses *Physcomitrium patens* (5 members), *Sphagnum fallax* (7 members), *Ceratodon purpureus* (3 members), the hornworts *Anthoceros agrestis Bonn* (one member)*, Anthoceros agrestis* Oxford (one member)*, Anthoceros punctatus* (one member), the lycophyte *Selaginella moellendorffii* (two members), the basal angiosperm *Amborella trichopoda* (nine members), and the 17 *Cr*RLK1Ls of *A. thaliana* (Fig. 1B). The *M. polymorpha Cr*RLK1L member grouped together with the *Cr*RLK1Ls of basal land plants and was in the same clade as, e.g., AtFER, AtHERK2, and AtANX1/2, while AtTHE1 was in a different one (Fig. 1B). Based on this phylogenetic position and the fact that the malectin-like domain of the *M. polymorpha Cr*RLK1L shared the highest amino acid identity with AtFER (Fig. S1A), we named it MpFER [48], [49], [60].

**Figure 1.**
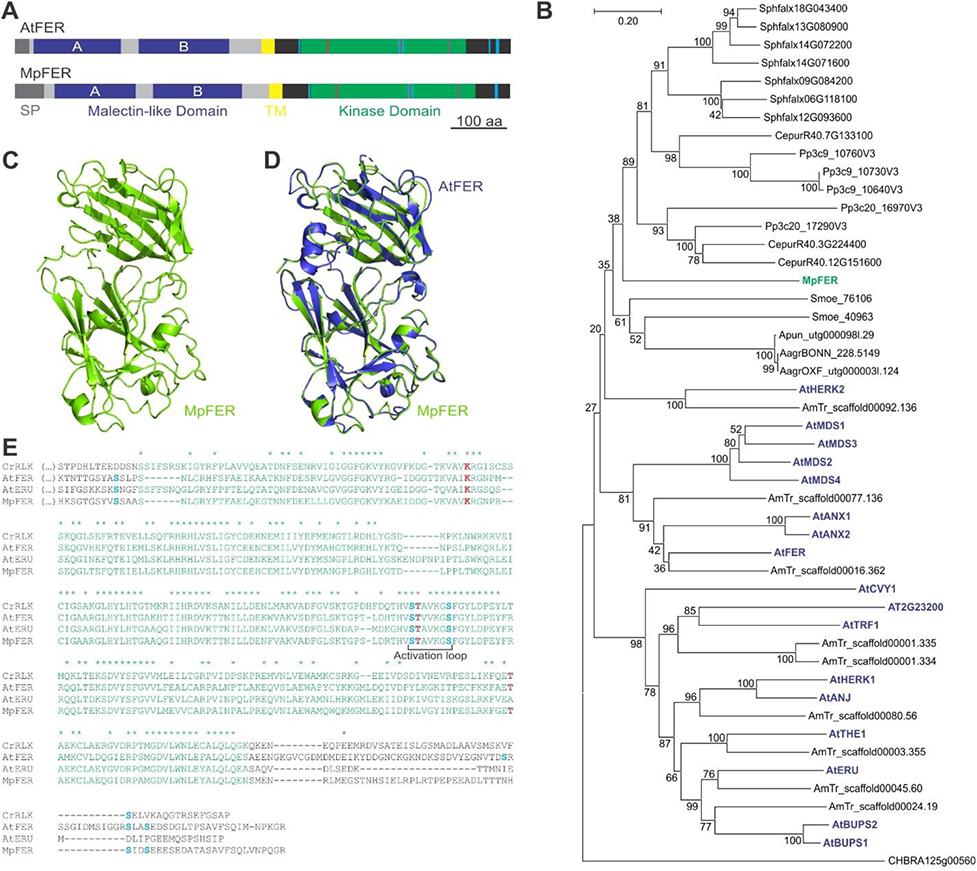
*Cr*RLK1Ls Are Conserved among Land Plants. (A) Representation of AtFER and MpFER proteins. The malectin domains are represented in blue and kinase domains in green. SP, signal peptide; TM, transmembrane domain (yellow). Amino acids important for regulation and activity are marked in red, violet, and light blue as described in Fig. 1E. (B) A rooted neighbor-joining tree of the amino acid sequence of the predicted malectin-like domain (corresponding to aa 76-431 of MpFER). *Cr*RLK1L members from *Marchantia polymorpha* (Mp, green), *Physcomitrium patens* (Pp), *Sphagnum fallax* (Sphfalx), *Ceratodon purpureus* (Cpur)*, Anthoceros agrestis* Bonn (AagrBONN)*, Anthoceros agrestis* Oxford (AagrOXR)*, Anthoceros punctatus* (Apun)*, Selaginella moellendorffii* (Smo), *Amborella trichopoda* (AmTr), and *Arabidopsis thaliana* (At, blue) were used. The best hit from *Chara braunii* (CHBRA) was also included. The numbers indicate the bootstrap values (%) from 1000 replications. The given scale represents a substitution frequency of 0.1 amino acids per site. (C) Cartoon representation of the predicted three-dimensional structure of the MpFER malectin-like domain, showing predicted alpha-helix and beta-sheet structures. (D) Structural superposition of the malectin-like domains of AtFER (blue) and MpFER (green). (E) Alignment of the cytoplasmic domains of MpFER, AtERU, AtFER, and *Cr*RLK. Kinase domains are in green, putative phosphorylation sites in light blue (Ser) and violet (Thr), and the conserved catalytic Lys in red. See also Fig. S1

Interestingly, no significant similarity to the MpFER malectin-like domain was found when compared to available genomic data of six Chlorophyte algae (*Chlamydomonas reinhardtii, Dunaliella salina, Volvox carteri, Coccomyxa subellipsoidea, Micromonas pusilla*, *Ostreococcus lucimarinus*) and expression data of six Charophycean algae (*Klebsormidium nitens, Nitella mirabilis, Coleochaete orbicularis, Spirogyra pratensis, Mesostigma viride*, *Closterium peracerosum–strigosum–littorale* complex). In *Closterium* complex, an RK named CpRLK1 expressed during sexual reproduction was suggested to be a *Cr*RLK1L homolog [62]. However, comparing the full sequence or the ECD of CpRLK1 with the predicted *M. polymorpha* proteome, higher identity to another RK was found (Mp1g17720 [Mapoly0001s0111]) (Figs S1B,C). Moreover, no RALF orthologs have been reported in algae, suggesting the origin of the *Cr*RLK1L pathway in land plants [48]. A similar conclusion was recently reached by analyzing the RALF family in *P. patens* [63]. Moreover, the best candidate obtained by BLAST analysis using the MpFER malectin-like domain against the genome of the Charophycean alga *Chara braunii* clustered outside the *Cr*RLK1L phylogenetic tree (Figs. 1B, S1B).

To further support MpFER as a member of the *Cr*RLK1L subfamily, the three-dimensional structure of its ECD was modelled [64] using the structure of the AtFER ECD (PDB 6a5b.1A, [36]) as a template (Fig. 1C). Overall, the predicted structure of the MpFER ECD is highly similar to that of AtFER, keeping an almost identical number of secondary structures (7 alpha-helices and 33 beta-sheets in the MpFER ECD [36] and 8 alpha-helix and 34 beta-sheets in the AtFER ECD, respectively), and a similar spatial disposition for 32 of the beta-sheet structures and 6 of the alpha-helices (Fig. 1C). When comparing the structural superposition of the AtFER and MpFER ECDs (Figs. 1D and S1D), we observed that most conserved parts of the malectin-like domain are in the core of the protein, while the more variable parts are located peripherally. Moreover, the 3D structure of the CpRLK1 ECD could not be properly modelled using the AtFER ECD (Fig. S1D,E), neither the one from *C. braunii* (Fig. S1D,F). Taken together, these results suggest that the malectin-like domain of MpFER could interact with similar proteins as AtFER does in *A. thaliana* and may thus be involved in similar pathways. Sequence comparison of the MpFER kinase domain showed a conservation of the activation loop and a Lys important for catalytic activity (Fig. 1E). Furthermore, several determined phosphorylation sites found in AtFER, AtERU, and CrRLK1 were conserved in MpFER (Fig. 1E) [14], [34], [65], [66]. Given that the genomes of *M. polymorpha* and *Anthoceros* spp. encode a single *Cr*RLK1L and this RK family is not present in Charophycean algae, we propose MpFER to be basal and orthologous to all other land plant *Cr*RLK1Ls, and that the *Cr*RLK1L family arose as plants conquered land.

### Mp*FER* is broadly expressed throughout the *M. polymorpha* life cycle

In *A. thaliana*, the expression patterns of the 17 *Cr*RLK1L members span vegetative and reproductive stages and both generations of the life cycle (reviewed in [44]), pointing to the importance of *Cr*RLK1Ls for basic cellular functions. To assess whether this pattern is reflected in the land plant with the most ancestral characteristics, a promoter fragment of 3.2 kb (*_pro_*Mp*FER*) was used to drive expression of a triple yellow fluorescent protein (trpVNS) with a nuclear localization signal (*_pro_*Mp*FER*:*trpVNS*) and transformed into *M. polymorpha* sporelings. All transformants expressing the trpVNS fluorescent protein (45 of 48 independent lines) exhibited an indistinguishable expression pattern (Fig. 2). During vegetative stages of gametophyte development, expression was observed in most cells of the thallus (Fig. 2A). In concordance with the importance of Mp*FER* for rhizoids [59], high trpVNS expression was observed in these tip-growing cells (Fig. 2B). While trpVNS was expressed in gemma cups, no expression was observed in mature gemmae, possibly due to their dormancy (Fig. 2C).

**Figure 2.**
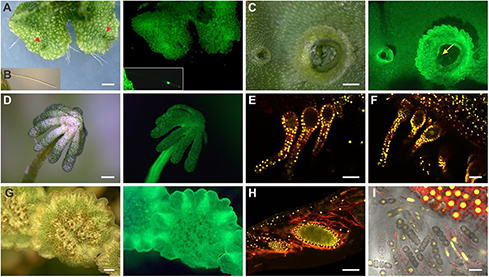
Mp*FER* Is Broadly Expressed in most Tissues throughout the *M. polymorpha* Life Cycle. Expression of *_pro_*Mp*FER*:*trpVNS* in different organs of male and female *M. polymorpha* plants. Fluorescence is visualized as either green (epifluorescence microscope) or yellow (confocal microscope) color, depending on the panel. (A to D) Bright field (left) and epifluorescence (right) images of a thallus, (A) rhizoid (C), gemmae cups (C), and an archegoniophore (D). Red arrows indicate meristematic zones, the yellow arrow gemmae. Scale bars, 0.5 mm. (E and F) Confocal images of archegonia before (E) and 2 days after fertilization (F). Scale bar, 50 μm. (G) Bright field (left) and epifluorescence (right) images of an antheridiophore. Scale bar, 1 mm. (H) Confocal image of antheridia. Scale bar, 100 μm. (I) Confocal image of sporophyte and spores. Scale bar, 25μm.

In mature female sexual organs, strong expression was observed in the entire archegoniophore (Fig. 2D). The archegonia expressed trpVNS in most cells, except for the egg cell (Fig. 2E). However, after fertilization, _pro_Mp*FER*:*trpVNS* became active in the zygote (Fig. 2F). In male sexual organs, trpVNS was broadly detected in the antheridial splash platform (Fig. 2G) and, specifically, in the spermatogenous tissue of the antheridia and the non-reproductive jacket cells surrounding them at different developmental stages (Fig. 2H). Inside the sporogonium, the sporogenous cells and subsequently the developing spores, but not the elater cells, strongly expressed trpVNS (Fig. 2I).

Taken together, Mp*FER* is expressed in most tissues during vegetative stages of gametophyte development as well as in the antheridia and archegonia during the sexual reproduction. Interestingly, the _pro_Mp*FER* promoter is not active in mature gemmae and the unfertilized, quiescent egg cell, although expression is initiated after dormancy and in the zygote. In concordance with the collectively ubiquitous expression of *Cr*RLK1Ls in *A. thaliana*, the Mp*FER* expression pattern reinforces the importance of this family for basic cellular functions in land plants.

### Mp*FER* controls cell expansion during vegetative gametophyte development

To analyze the function of Mp*FER*, three independent artificial microRNA constructs targeting Mp*FER* were designed (amiR-Mp*FER*), based on the endogenous microRNA Mp*miR160* (Fig. S2, [67]) and driven by the ubiquitous *_pro_*Mp*EF1* promoter [68]. While amiR-Mp*FER*1 and amiR-Mp*FER*2 target sites in the ECD coding sequence, amiR-Mp*FER*3 targets the sequence encoding the kinase domain (Fig. 3A,B). For each amiR-Mp*FER* construct, several independent transgenic lines were obtained that showed a reduction in thallus size (Figs. 3C,D and S3A). Mp*FER* levels were quantified by qRT-PCR in two independent amiR-Mp*FER*2 and amiR-Mp*FER*3 lines each, which showed a similar phenotype (Fig. 3E). Mp*FER* expression ranged from 10% to 20% of the wild-type level in all amiR-Mp*FER* lines analyzed (Fig. 3E).

**Figure 3.**
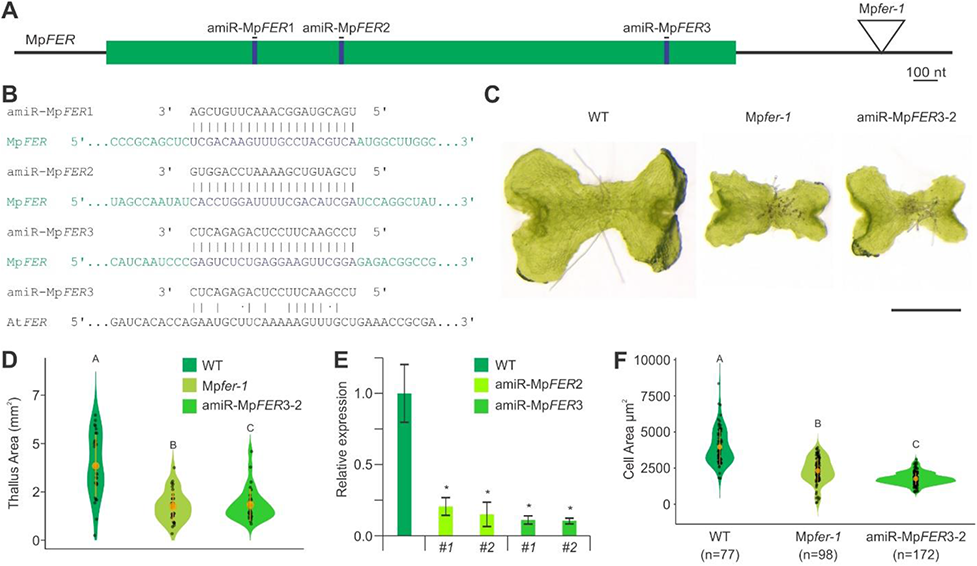
Mp*FER* Controls Cell Size during Gametophyte Development. (A) Representation of the mature Mp*FER* mRNA. The coding sequence is in green, locations of the amiRNA target sites in blue. The location of the T-DNA insertion in Mp*fer-1* is shown. (B) Base complementary of mature amiR-Mp*FER*1, amiR-Mp*FER*2, and amiR-Mp*FER*3 with the Mp*FER* transcript. Very low complementary was found between amiR-Mp*FER3* and At*FER*. (C) Representative pictures of 10-day old gemmalings of wild-type (WT), Mp*fer-1*, and amiR-Mp*FER*3-2 lines. Scale bar, 1 mm. (D) Violin plot of thallus area of 14-day old gemmalings of WT, Mp*fer-1*, and amiR-Mp*FER*3-2 lines. n = 30. Different letters indicate significant differences according to the non-parametric Kruskal-Wallis test and Dunn’s test for multiple comparisons (P < 0.001). Orange circles indicate the group mean and the corresponding vertical bars the standard deviation for each group. (E) Relative expression level of Mp*FER* in thallus tissue from WT and two independent insertion lines of amiR-Mp*FER*2 and amiR-Mp*FER*3, as measured by qRT-PCR. Mp*EF1* was used as internal control. Shown are means ± standard errors of the means (SEM) of three biological replicates. Statistical analysis was performed by one-way analysis of variance (ANOVA) followed by a post-hoc Duncan test (*P < 0.01). (F) Violin plot of cell size in WT, *Mpfer-1*, and amiR-Mp*FER*3-2 lines. Only areas with fully expanded cells in approximately 3-week old plants were used for measurements. Different letters indicate significant differences according to the non-parametric Kruskal-Wallis test and Dunn’s test for multiple comparisons (P < 0.001). Orange circles indicate the group mean and the corresponding vertical bars the standard deviation for each group. See also Figs. S2,S3

Previously, the T-DNA insertion mutant Mp*fer-1* was identified in a screen for mutants with defective rhizoids ([59], referred to as Mp*the*). As the T-DNA inserted into the 3’ UTR of the Mp*FER* gene (Fig. 3A; [59]), Mp*FER* expression was analyzed by qRT-PCR but it showed no reduction compared to the wild type (Fig. S3B), suggesting that the T-DNA insertion does not affect Mp*FER* transcription. Sequencing of the Mp*FER* coding sequence in the Mp*fer-1* mutant did not show any mutation, while 3’ RACE-PCR showed that transcript terminated about 1.3 kb inside the T-DNA and was not polyadenylated (Fig. S3C). Because thallus size of Mp*fer-1* mutants is similar to that of amiR-Mp*FER* lines (Figs. 3C,D and S3A) and efficient translation depends on the presence of a poly(A)-tail [69], we propose that MpFER protein levels are reduced in the Mp*fer-1* mutant despite normal transcript levels.

Rhizoids are tip-growing cells analogous to the root hairs in flowering plants (angiosperms). Reduced Mp*FER* activity strongly impaired rhizoid formation and the rhizoids collapsed (Fig. S3D). This had also been reported for the Mp*fer-1* mutant [59] and is in concordance with the function of At*FER* and At*ANX1/2* in tip-growing root hairs [13], [17], [34], [70] and pollen tubes [9], [15], respectively. As *Cr*RLK1L members are important for cell expansion in angiosperms [8], [10], [12], [20], [35], [71], [72], cell size of epidermal thallus cells in fully differentiated regions with minimal growth was measured in Mp*fer-1* and amiR-Mp*FER* lines, showing a reduction in cell area as compared to the wild type (Figs. 3F and S3E,F). The average epidermal cell area was similarly reduced, although the distribution of cell areas differed between amiR-Mp*FER* lines and Mp*fer-1* (Figs. 3F and S3E,F).

In conclusion, Mp*FER* has a fundamental role in rhizoid formation and growth during vegetative development. These results suggest an ancestral and conserved function of the *Cr*RLK1Ls in polar cell growth and cell expansion.

### Mp*FER* is important for the morphological integrity of the gametophyte

Given that Mp*fer-1* and amiR-Mp*FER* lines retain some MpFER activity, we generated Mp*fer* knock-out mutants with the goal to unveil potentially hidden *Cr*RLK1L functions by generating a plant without any *Cr*RLK1L activity. Using CRISPR/Cas9, two sites in the ECD-coding sequence of Mp*FER* were targeted (Fig. 4A,B). At least 10 independent lines were selected and analyzed for each target site. All plants with severely affected thallus development contained a frame shift mutation at the respective target site (Fig. 4B,C) while Mp*FER* was not affected in transgenic plants with normal development.

**Figure 4.**
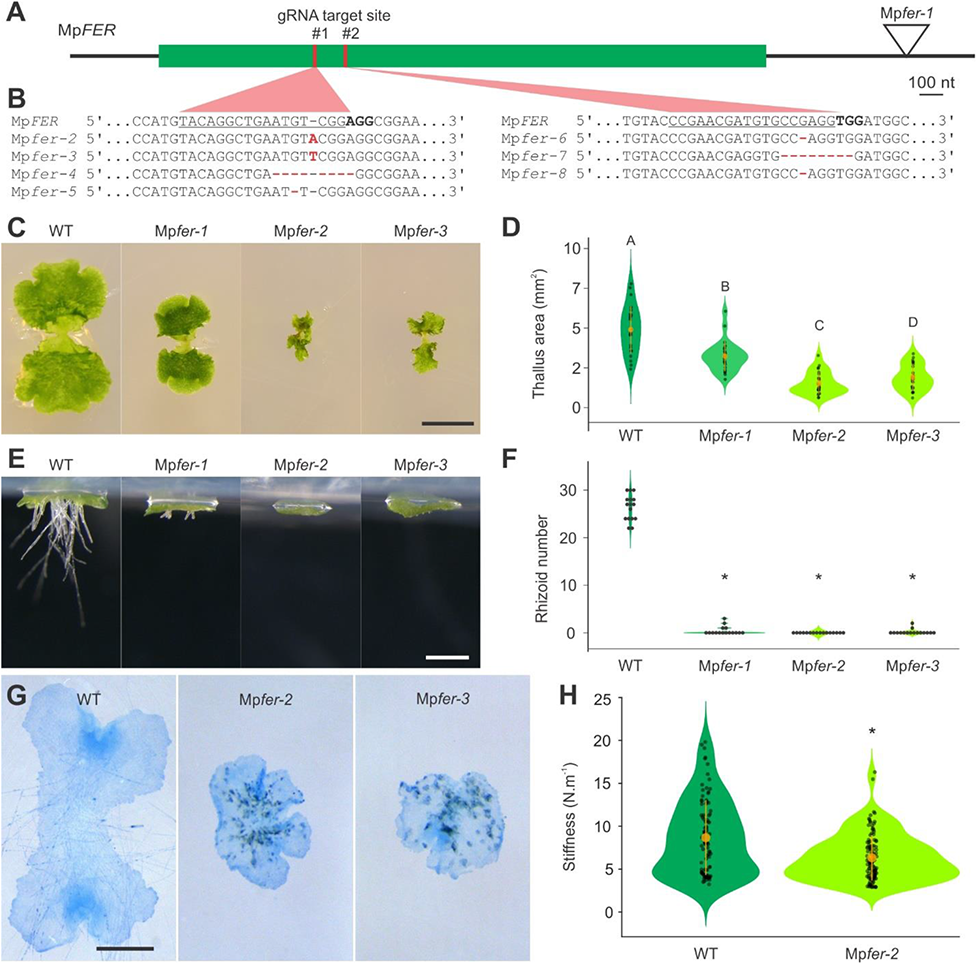
The Integrity of Thalli from Mp*fer* Knock-out Lines Is Severely Affected. (A) Representation of the mature Mp*FER* mRNA with the coding sequence in green, and the location of the gRNA target sites in red. (B) Sequences of gRNA target sites in the confirmed Mp*fer* knock-out mutants. Deletions and insertions are highlighted in red. (C) Representative pictures of 14-day old gemmalings of the wild type (WT), Mp*fer-1*, and two independent Mp*fer* knock-out mutants. Scale bar, 0.5 cm. (D) Violin plot of thallus area of 7-day old gemmalings of WT, Mp*fer-1*, and two independent Mp*fer* knock-out mutants. n = 25. Different letters indicate significant differences in a one-way analysis of variance (ANOVA) follow by a post-hoc Duncan test (P < 0.01). Orange circles indicate the group mean and the corresponding vertical bars the standard deviation for each group. (E) Representative pictures of 7-day old gemmalings growing in upside-down plates of WT, Mp*fer-1*, and two independent Mp*fer* knock-out mutants. Scale bar, 500 µm. (F) Violin plot of rhizoid number in 3-day old gemmalings of WT, Mp*fer-1*, and two independent Mp*fer* knock-out mutants. n = 16. Statistical analysis was performed by one-way analysis of variance (ANOVA) followed by a post-hoc Duncan test (*P < 0.01). (G) Trypan blue staining of 7-day old gemmalings of WT, Mp*fer-1*, and two independent Mp*fer* knock-out mutants. Scale bar, 1 mm. (H) Violin plot of the reduction in the apparent stiffness of gemmaling tissue in Mp*fer-2* compared to the WT. Statistical analysis was performed using a Kruskal-Wallis test followed by a Tukey-Kramer post-hoc test (*P < 0.001). Orange circles indicate the group mean and the corresponding vertical bars the standard deviation for each group.

The thallus area of the newly generated, amorphic knock-out mutants was more strongly reduced compared to Mp*fer-1* [59], confirming that, like the amiR-Mp*FER* lines, Mp*fer-1* is a hypomorphic mutant (Fig. 4C,D). In contrast, rhizoid integrity was similarly affected in hypo- and amorphic Mp*fer* alleles, suggesting that rhizoid formation is more sensitive to reduced Mp*FER* activity than thallus growth (Fig. 4E,F). Despite of the strong impact on thallus development, both hypo- and amorphic Mp*fer* mutants form gemma cups.

The strong disruption of thallus growth in amorphic Mp*fer* mutants prompted us to investigate whether cells of the thallus died in the absence of Mp*FER* activity. Indeed, using tryphan blue staining, we detected dead cells in these mutants, indicating a loss of morphological integrity (Fig. 4G). To investigate the mechanical properties of thallus cells, we measured their apparent stiffness using cellular force microscopy on 3-day old gemmalings and found that amorphic Mp*fer-2* mutants had a lower apparent stiffness than the wild type (Fig. 4H). The apparent stiffness of the thalli depends on both the stiffness of the cell wall and the turgor pressure of the cells. Since the apparent stiffness is lower in Mp*fer-2* mutants, it is likely that a reduction in the stiffness their cell wall causes the cells to burst and die.

### Some reproductive but not all CWI sensing functions of *Cr*RLK1Ls are conserved in land plants

In *A. thaliana*, at least seven *Cr*RLK1L family members play a role in reproduction, with At*ANX1/2* and At*BUPS1/2* being important for pollen tube growth [9], [15], [19], [33] and At*FER*, At*HERK1*, and At*ANJ* for pollen tube reception by the synergid cells [7], [18]. To characterize the function of Mp*FER* in reproductive development, amiR-Mp*FER3* lines were transferred to sexual organ-inducing conditions [73]. The number of antheridiophores produced was significantly less in amiR-Mp*FER* lines compared to the wild type (Figs. 5A and S3G) and the antheridiophore splash platforms were also smaller (Fig. 5B). The reduction in Mp*FER* expression in antheridiophores of amiR-Mp*FER* lines was confirmed by qRT-PCR and results were comparable to those using thallus tissue (Fig. 5C). A pronounced reduction in spermatogenous tissue was observed in many antheridia isolated from amiR-Mp*FER*3-3 lines (Fig. 5D,E). Whether this phenotype is caused by reduced cell expansion as was observed in thalli or by a lower cell division rate is currently unclear. Moreover, although the plants were grown under optimal conditions in axenic culture boxes, the effects on reproductive structures could be partly indirect due to the reduction in plant size and/or rhizoid function.

**Figure 5.**
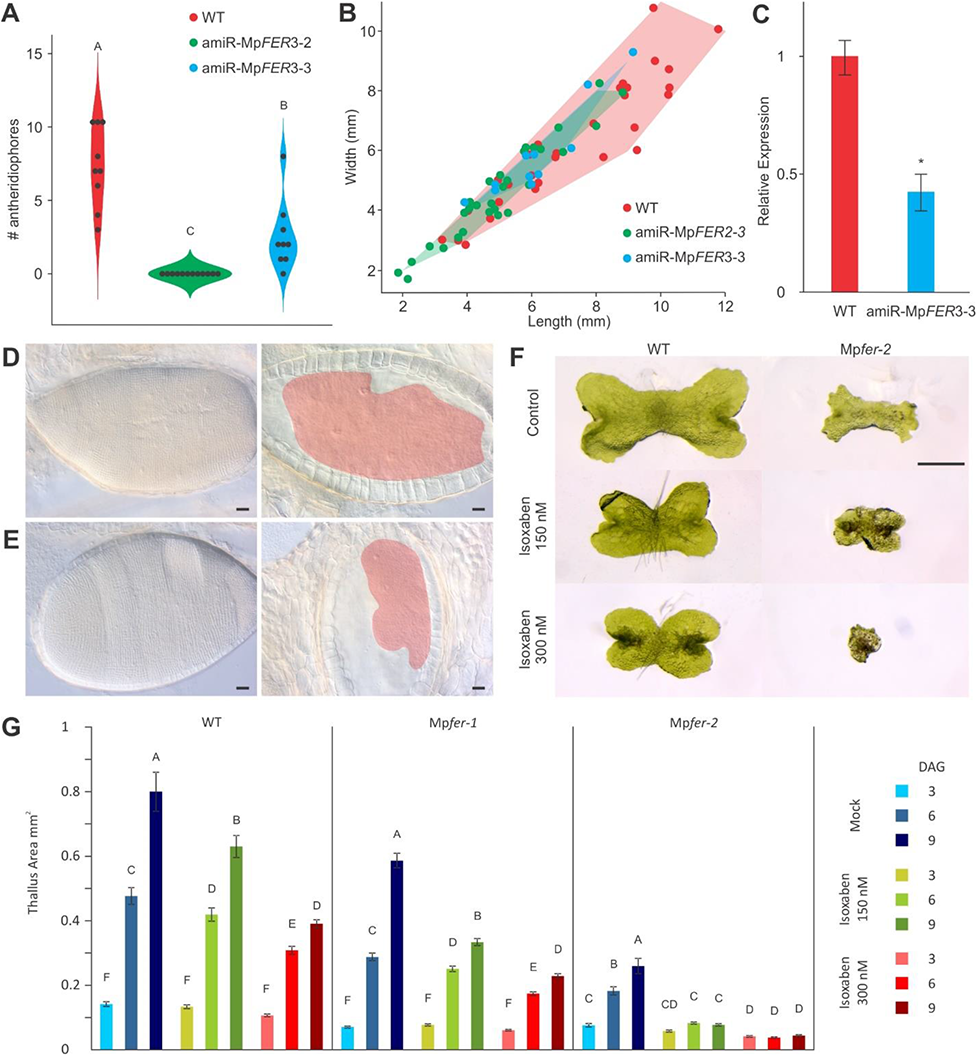
Conservation of *Cr*RLK1L Function in Land Plants. (A) Violin plot of the number of antheridiophores produced per plant for wild-type (WT, n=9) and two independent amiR-Mp*FER*3 lines (amiR-Mp*FER*3-2, n = 9; amiR-Mp*FER*3-3, n = 12). Different letters indicate a significant difference in a one-way analysis of variance (ANOVA) followed by a post-hoc Duncan test (P < 0.01). (B) Antheridiophore splash platform size distribution of WT (n = 32), amiR-Mp*FER*2-3 (n = 34), and amiR-Mp*FER*3-3 (n = 11) lines. For all antheridia with a stalk >8 mm, length and width of the platform were recorded. (C) qRT-PCR of Mp*FER* levels in antheridiophores in a WT and amiR-Mp*FER*3-3 line. Mp*EF1* was used as internal control. Shown are means ± SEM of three biological replicates. Statistical analysis was performed by one-way analysis of variance (ANOVA) followed by a post-hoc Duncan test (*P < 0.01). (D and E) Mature antheridia of a WT (D) and amiR-Mp*FER*3-3 line (E). Bright field images (left) and in cross-section (right). The spermatogenous areas are indicated in red. Scale bar, 100*μ*m. (F) Representative pictures of 6-day old gemmalings of WT and Mp*fer-2* mutants at different isoxaben concentrations. Scale bar, 1 mm. (G) Thallus area of gemmalings of WT, Mp*fer-1*, and Mp*fer-2* lines growing on media containing different isoxaben concentrations. n = 35. Shown are means ± SEM of two biological replicates. Different letters indicate a significantly difference in a one-way analysis of variance (ANOVA) followed by a post-hoc Duncan test (P < 0.05). See also Fig. S3

To determine male fertility of these amiR-Mp*FER* lines with reduced spermatogenous tissue, we performed crosses with wild-type female plants. The spermatocyte concentration harvested from wild-type antheridia was almost 3-fold higher than from amiR-Mp*FER* lines and was adjusted using a hemocytometer. In just one of the crosses (n=20), a single sporophyte was formed, resulting in 0.05 ± 0.05 (mean ± SE) sporophytes per archegoniophore, whereas the same females crossed with wild-type males yielded 9.95 ± 2.33 (mean ± SE) sporophytes per archegoniophore. In summary, Mp*FER* plays a role in antheridiophore development and spermatocytes of plants with reduced Mp*FER* activity are poorly fertile. Thus, *Cr*RLK1Ls have a conserved role in reproduction in addition to their role in cell expansion and integrity during vegetative development. Some *A. thaliana Cr*RLK1Ls are involved in surveying CWI and in inhibiting growth when CWI is impaired. Cell wall defects caused either by mutations affecting cellulose biosynthesis or by treatment with isoxaben, a cellulose synthesis inhibitor, can be suppressed by a mutation in At*THE1* [8], [29]. To investigate whether Mp*FER* also acts as CWI sensor, we treated wild-type, Mp*fer-1*, and Mp*fer-2* gemmae with isoxaben (Fig. 5F,G). As in the wild type, isoxaben affected the development of gemmae with reduced Mp*FER* activity, all growing progressively less with increasing isoxaben concentration (Fig. 5F,G). Moreover, isoxaben treatment was lethal for Mp*fer*-2 knock out mutants (Fig. 5F,G). Similar observations were reported for At*fer-4*, where isoxaben treatment increased the number of dead cells in hypocotyls [74]. These results suggest that the function of At*THE1* in repressing cellular growth when CWI is impaired appeared later in the course of land plant evolution or was lost in the *Marchantia* lineage.

### The *Cr*RLK1L signaling pathway is conserved in land plants

As downregulation of Mp*FER* or At*FER* produce similar phenotypes concerning polarized growth and cell expansion in *M. polymorpha* and *A. thaliana*, respectively, interspecific complementation was attempted. First, the coding sequence of Mp*FER* fused to the Green Fluorescent Protein (GFP) gene under the control of the viral *35S* promoter (*_pro_35S*:Mp*FER-GFP*) was transformed into heterozygous At*fer-1* mutants. Then, GFP expression at the cell periphery and complementation of the bursting root hair and reduced rosette size phenotypes were assessed in transgenic plants (Fig. 6A). At*fer-1* homozygotes expressing the MpFER-GFP protein look similar to At*fer-1* mutants, indicating that Mp*FER* does not complement these At*fer-1* loss-of-function phenotypes (Fig. 6A). While we cannot exclude that this is because we used a heterologous promoter, we consider this unlikely as expression the *35S* promoter is routinely used to complement root hair phenotypes [75], [76]. In a complementary experiment, the coding sequence of At*FER* fused to the Citrine gene under the control of the Mp*EF1* promoter (*_pro_*Mp*EF1:*At*FER-Cit*) was transformed into amiR-Mp*FER*3-2 plants. Ten independent lines with Citrine expression at the cell periphery were phenotypically characterized (Fig. 6B,C). All had bursting rhizoids and a reduced thallus size, similar to the parental amiR-Mp*FER*3-2 line, suggesting that At*FER* does not rescue the vegetative phenotypes produced by down-regulation of Mp*FER*. A sequence alignment of At*FER* with amiR-Mp*FER3* showed not complementarity (Fig. 3B), confirming that the failure to complement is not cause by a potential downregulation of At*FER* by amiR-Mp*FER3*. Taken together, these results show that MpFER and AtFER cannot substitute each other with respect to their function in cell expansion and tip growth.

**Figure 6.**
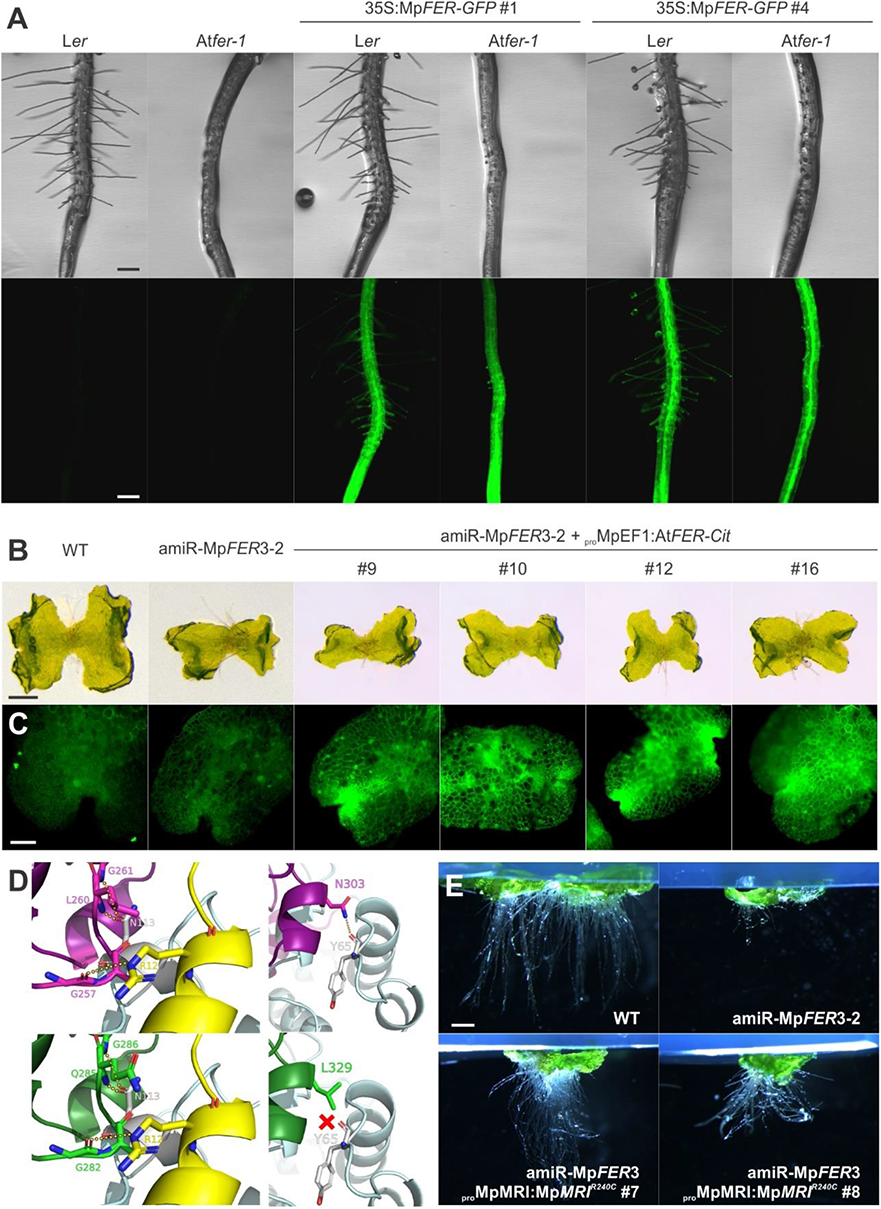
Interspecific Complementation of *A. thaliana* and *M. polymorpha* Plants with Reduced Levels of *FER* Activity. (A) Roots of independent *A. thaliana* lines expressing 35S:Mp*FER-GFP* in L*er* wild-type (WT) and At*fer-1* plants. Scale bar, 200 μm. (B) 10-day old gemmalings of WT and amiR-Mp*FER*3-2 *M. polymorpha* plants with and without expression of At*FER-Cit*. Scale bar, 0.5 mm. (C) Citrine expression in 7-day old gemmalings of WT and amiR-Mp*FER*3-2 plants with and without expression of At*FER-Cit*. Scale bar, 100 μm. (D) Cartoon representation of the *Cr*RLK1L^ECD^ polar interactions sites with LRE/RALF when forming a protein complex. Upper panels correspond to contact areas between AtFER (purple), AtLLG2 (blue-grey), and AtRALF23 (yellow), and lower panels to the modeled MpFER structure (green) superimposed onto the same complex. Polar interactions are depicted as doted yellow lines. Left panels show the predicted polar interactions of the complex that are conserved between *A. thaliana* and *M. polymorpha*, whereas the right pannels show the polar interaction between AtFER and AtLLG2 that is predicted to be absent between MpFER and AtLLG2. (E) Suppression of the amiR-Mp*FER* phenotype by a dominant-active version of Mp*MRI* under its own promoter (*_pro_*Mp*MRI*:*MRI^R240C^*). Scale bar, 1 mm. See also Figs. S4-S6

In *A. thaliana*, it was shown that AtFER forms a complex with AtRALF peptides and the LRE/LLG co-receptors [36]. As no interspecific complementation was observed, the conservation of binding surfaces between *Cr*RLK1L, RALF, and LRE homologs were analyzed. The *M. polymorpha* genome encodes three RALF peptides (MpRALF: Mp1g27120 [Mapoly0002s0166]; MpRALF2: Mp2g21670 [Mapoly0040s0047]; MpRALF3: Mp7g07270 [Mapoly0076s0067]) [47], [48]. We performed a phylogenetic analysis using the amino acid sequences of the predicted mature peptides of *A. thaliana* and *M. polymorpha* RALFs (Fig. S4A). All MpRALF peptides clustered together with AtRALFs known to interact with *Cr*RLK1L receptors, including AtRALF1 and AtRALF23, which serve as AtFER ligands [23], [34] (Fig. S4A). The MpRALFs shared the four conserved Cys residues as well as the YXXY and YY motifs with the AtRALF1 subgroup (Fig. S4B). However, only MpRALF1 and MpRALF3 had an RRXL motif important for S1P cleavage (Fig. S4B,C)[77], consistent with the presence of one S1P ortholog in basal plants (Mp8g07990 [Mapoly0155s0018], Fig. S4D). There are two MpLRE/LLG proteins encoded in the *M. polymorpha,* genome (Fig. S5A,B). Both the MpLRE1 (Mp5g09600 [Mapoly0048s0110]) and MpLRE2 (Mp4g22100 [Mapoly0090s0020]) proteins showed conservation of the eight Cys and the ND motif distinctive of the family (Fig. S5B); however, MpLRE1 did not contain a GPI anchoring site (Fig. S5B). Structure prediction using AtLLG1 as a template [36] showed a conserved general structure (Fig. S5C,D). Some of the Mp*RALF* and Mp*LRE* family members show a similar expression pattern as Mp*FER* (Fig. S5F), suggesting that the corresponding proteins could form a complex similar to that described in *A. thaliana*.

To gain insights into the formation of a putative *Cr*RLK1L signaling complex, the MpFER/MpRALF/MpLRE complex was modelled. The general structure appeared conserved (Fig. S5E); however, analysis of conserved sites between *M. polymorpha* and *A. thaliana* showed a low amino acid conservation in the interacting surfaces of the complex subunits (Fig. 6D). This suggests that the lack of interspecific complementation may be due to differences in amino acids that are important for complex formation, and that the proteins forming the complex have diverged, following distinct routes of co-evolution in the two lineages.

Because interspecific complementation of the amiR-Mp*FER* phenotypes was unsuccessful, we asked whether suppression using a downstream component of *Cr*RLK1L signaling identified in *A. thaliana* was possible. AtMRI, a receptor-like cytoplasmic kinase, acts downstream of At*FER* or At*ANX1/2* in the regulation of polar tip growth [78], [79]. The dominant AtMRI^R240C^ mutation can suppress pollen tube rupture in the At*anx1*/At*anx2* double mutant and the root hair defects of At*fer-4* [78], [79]. A mutation in Mp*MRI* (aka Mp*PTI*) with a similar bursting rhizoid phenotype as observed in Mp*fer-1*, was identified in the same genetic screen for *M. polymorpha* mutants with defective rhizoids [59]. To test whether Mp*MRI* is a conserved downstream component of the Mp*FER* signal transduction pathway, a dominant-active form of the protein equivalent to AtMRI^R240C^ was transformed into the amiR-Mp*FER*3-2 line, driven by 2 kb of the endogenous promoter (*_pro_*Mp*MRI:*Mp*MRI^R240C^*). Several independent lines showed a partial restoration of rhizoid growth; however, we also observed defects in thallus development of lines with higher levels of Mp*MRI* expression, mainly abnormalities in the epidermis (Figs. 6E and S6A-E). This suggest that Mp*MRI* acts downstream of Mp*FER* in the signal transduction pathway controlling polarized growth since the origin of land plants, in agreement with a recent report on the functional characterization of Mp*MRI* [60].

### Overexpression of Mp*FER* affects morphological integrity

Because overexpressing At*FER* in *M. polymorpha* did neither lead to any obvious phenotypes nor complementation of amiR-Mp*FER* lines (Fig. 6B,C), we also overexpressed At*FER* and Mp*FER* in the wild type. As in the amiR-Mp*FER* background, overexpression of AtFER (*_pro_*Mp*EF1:*At*FER-Cit*) in a wild-type background had no effect, consistent with the idea that At*FER* is unable to form a complex with the corresponding *M. polymorpha* proteins. However, when expressing an MpFER-Citrine fusion protein (*_pro_*Mp*EF1:*Mp*FER-Cit*) in wild-type plants, thallus development was strongly affected (Fig. 7A). As expected, MpFER-Cit localized to the membrane (Fig. 7A inset), and higher protein levels correlated with more severe phenotype (Fig. S6F,G). Scanning electron microscopy of epidermis of plant expressing high levels of Mp*FER* showed defects in the formation of air chambers and air pores (Fig. 7B), similar to what we observed when expressing the dominant-active Mp*MRI^R240C^* (Fig. S6A-C). This phenotype is reminiscent of the *quasimodo1* mutant in *A. thaliana*, in which cell walls have a reduced pectin content and cell adhesion is compromised [80]. Moreover, overexpression of Mp*FER* affects the number of lobes in the antheridial receptacle, producing only 4 instead of 8 (Fig. S6H), supporting the importance of Mp*FER* during sexual development. Considering the importance of Mp*FER* for normal rhizoid formation, we also analyzed rhizoids in the *_pro_*Mp*EF1:*Mp*FER-Cit* lines: while rhizoid morphology looked normal, a decrease in rhizoid number was observed (Fig. 7C,D). To analyze cell wall properties of *MpFER* overexpressor lines, chlorophyll efflux rates were measured, as an indicator for epidermal permeability (Fig. 7E). Overexpression of *MpFER* clearly reduced the chlorophyll efflux compared to the wild type, while the opposite was observed in Mp*fer-2* mutants (Fig. 7E). Taken together, the results obtained by overexpressing Mp*FER* support a role of Mp*FER* in cell wall formation and the maintenance of tissue integrity during plant morphogenesis.

**Figure 7:**
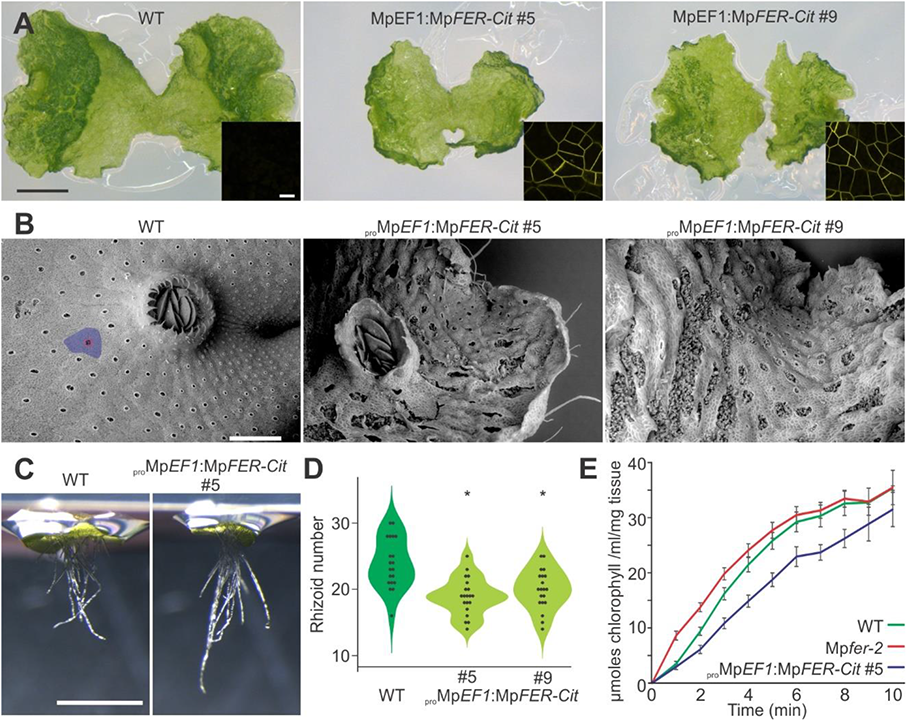
Overexpression of Mp*FER* Affects Morphological Integrity. (A) Representative pictures of 10-day old gemmalings of wild-type (WT) and two different lines overexpressing Mp*FER* (*_pro_*Mp*EF1:*Mp*FER-Cit)*. Scale bar, 1 mm. Inset: Citrine expression at the plasma membrane in *_pro_*Mp*EF1:*Mp*FER-Cit* lines *#5* and #9 as observed under confocal microscopy. Scale bar, 20 µm. (B) Scanning electron microscopical images of thalli from 20-day old plants of WT and *_pro_*Mp*EF1:*Mp*FER-Cit* lines *#5* and #9. The blue region demarks an air chamber, while air pore cells are colored in red. Scale bar, 500 µm. (C) Representative pictures of 3-day old gemmalings growing in upside-down plates of WT and *_pro_*Mp*EF1:*Mp*FER-Cit #5* lines. Scale bar, 1 mm. (D) Violin plot of the number of rhizoids in 3-day old gemmalings of WT and *_pro_*Mp*EF1:*Mp*FER-Cit* lines *#5* and #9. n = 20. Statistical analysis was performed by one-way analysis of variance (ANOVA) followed by a post-hoc Duncan test (*P < 0.01). (E) Graphs showing amount of effluxed chlorophyll as a function of time from 15-day old gemmalings of WT, Mp*fer-2* and *_pro_*Mp*EF1:*Mp*FER-Cit #5* lines. See also Fig. S6

## Discussion

During land plant evolution, many developmental aspects have changed in order to adapt to new environments, producing the enormous diversity observed in the plant kingdom. However, the control and maintenance of CWI remained a key aspect for the biology of a plant cell. CWI sensing is not only important for vegetative growth by cell expansion, but it is also essential for sexual reproduction and defense responses.

The *Cr*RLK1L family was first characterized for its importance during angiosperm fertilization, a process comprising aspects of polar cell elongation, CWI control, and cell-cell communication [7], [31], [32]. Lately, different members of this family were found to carry out diametrically opposite roles in diverse aspects of plant development, which has made it difficult to define the core or ancestral function of this gene family comprising 17-members in *A. thaliana* [7]–[13], [15]–[21], [33].

Here, we report the characterization of the single *Cr*RLK1L gene encoded in the genome of *M. polymorpha*. Structurally, the MpFER ECD and kinase domain share similarities with previous characterized *Cr*RLK1L members, including the malectin-like domain and conserved phosphorylation sites. Conservation of RALF and LRE members in *M. polymorpha* suggests that MpFER also forms a complex at the plasma membrane [36]. While modeling predicted that the general structure of the complex was conserved, the interaction surfaces seem to have diverged and co-evolved independently in the respective lineages.

We observed a broad involvement of Mp*FER* at both vegetative and reproductive stages of development: During vegetative growth of the gametophyte, Mp*FER* is required for rhizoid formation and cell expansion but it also plays a role in male gametogenesis and is expressed in female reproductive organs. These findings suggest that the *Cr*RLK1Ls have, also in ancestral land plants, held roles in various aspects of both vegetative and reproductive development. Thus, the role of *Cr*RLK1Ls in angiosperm reproduction does not represent a derived feature from a purely vegetative function in cell elongation in bryophytes but constitutes a conserved feature of this gene family.

Moreover, the almost ubiquitous expression of Mp*FER* in *M. polymorpha* is consistent with the collective expression of different *Cr*RLK1L members in essentially all tissues and organs of *A. thaliana* [44]. Thus, gene duplication allowed the expansion of the *Cr*RLK1L family in angiosperms, diversifying their expression patterns and functions but also leading to genetic redundancy [9], [18], [33], [44].

*Cr*RLK1L members are important sensors of CWI during polarized growth, both in pollen tubes [9], [33] and root hairs [17]. In early divergent land plants, rhizoids are tip-growing cell with a function analogous to that of root hairs, important for taking up nutrients and water [59], [81]. Based on the conservation of many genes controlling the development and growth of tip-growing cells forming rooting systems, it was suggested that these pathways were active in the earliest land plants that existed about 470 million years ago [59]. That Mp*FER* also controls rhizoid integrity points to the importance of the *Cr*RLK1L pathway for polarized growth since the origin of land plants. Moreover, the fact that amorphic Mp*fer* mutants contain many dead cells demonstrates the importance of the *Cr*RLK1L pathway for cellular integrity during vegetative growth. As the occurrence of dead cells was recently also reported in At*fer-4* mutant seedlings [74], the function of this pathway in maintaining cellular integrity may also be ancestral. Whether Mp*FER* also plays a central role in plant innate immunity, as it does in angiosperms [16], [23], remains to be determined.

The signaling mechanisms downstream of Mp*FER* in *M. polymorpha* development are still unclear. Mp*MRI*, the *M. polymorpha* ortholog of At*MRI*, which acts downstream of AtFER and AtANX1/2 in *A. thaliana* during polarized growth [78], does have a similar rhizoid phenotype as Mp*FER* [59], [60]. The suppression of the rhizoid phenotype in amiR-Mp*FER* transformants with the constitutively active form MpMRI^R240C^ corroborates a conserved *Cr*RLK1L signaling cascade during polarized growth in *M. polymorpha* but whether this pathway also relies on Ca^2+^ and reactive oxygen species (ROS) as second messengers as it does in *A. thaliana* remains to be determined.

In *A. thaliana*, loss-of-function mutations in the motor protein AtKINESIN-13a, a microtubule-based motor, cause cell elongation in petals, leaves, and hypocotyls concomitant with changes in cell wall composition in leaves [82]. This effect depends on functional AtTHE1 and does not occur in At*kinesin-13-a/*At*the1* double mutants. Interestingly, in this case, AtTHE1 stimulates cell elongation in response to defects in cell wall deposition [82]. Based on our results, we have no indication that Mp*FER* acts as a CWI sensor and regulates growth similar to At*THE1* in response to cell wall disturbances. We have not observed a context in *M. polymorpha* development in which Mp*FER* inhibits cell expansion. Therefore, the ability to restrict growth in response to disturbances in CWI could represent an evolutionary derived feature, as gene duplication allowed for the diversification and specialization of *Cr*RLK1Ls to regulate cellular growth in a more complex and context-dependent way. The capacity to flexibly regulate growth appears to be fundamentally context-dependent, as a similar discrepancy in promoting or inhibiting growth is also known for some well-studied growth promoting agents, like the phytohormones auxin and brassinosteroid, which can have growth-inhibitory effects dependent on tissue and concentration. The reverse was noted for the phytohormones ethylene and abscisic acid, which are usually considered growth inhibitors but can also promote growth, dependent on the context [83]–[85].

Taken together, our results suggest an ancestral and conserved function of the *Cr*RLK1Ls in polar cell growth and cell expansion, but also during sexual reproduction. In addition, we probe the relevance of *Cr*RLK1L members for cell integrity and morphogenesis. In angiosperms, *Cr*RLK1Ls occupy an important position at the interface of developmental and environmental inputs, which are integrated by a downstream signaling machinery that controls cell shape and polar growth to ensure normal development, e.g., by preventing cell rupture upon CWI defects. Whether the same downstream machinery is utilized in a similar fashion in *M. polymorpha* remains to be shown, even though the similarity in function points to a related mechanism. Even less is known about upstream aspects of *Cr*RLK1L signaling. Future studies on the transcriptional regulation and the upstream components, such as the putative RALF ligand(s), in the highly simplified *M. polymorpha* system could provide fundamental insights into the molecular mechanism of the *Cr*RLK1L pathway, which is so central to land plant physiology, growth, and development.

## Materials and methods

### Experimental model

*M. polymorpha* subsp. *ruderalis* (Bischl. & Boissel.-Dub.) plants of the Tak-1 accession were grown on sterile culture on half-strength Gamborg’s B5 basal medium (PhytoTechnology Laboratories), supplemented with 1% phytoagar in Petri dishes sealed with air-permeable tape. The plants were kept under fluorescent light and long-day conditions (16 h light at 22°C, 8 h dark at 20 °C) in Percival growth cabinets (models AR-41L3 and AR-41/L2). Alternatively, to induce sexual reproduction, plants were cultivated in jiffy pots filled with a 1:1 mixture of soil (“Einheitserde D73 + Bims”, Universalerde) and sand or in culture boxes in sterile culture on half-strength Gamborg’s B5 basal medium (Duchefa Eco2 Box, #E1654), under fluorescent light supplemented with far-red light (GreenPower LED module HF far-red, #929000464503 and #929000632103 Philips). Wild-type plants in all experiments were *Arabidopsis thaliana* L. (Heyn) accession L*er*-0. The At*fer-2* mutant was obtained from the Signal Collection at the Salk Institute. Plants were grown in soil (ED73; Universalerde), covered with a thin quartz sand layer, under long photoperiods (16 hs light/8 hs dark) at 23°C and 60% relative humidity.

### Phylogenetic analysis

Protein sequences were identified in plant genomes via BLASTp searches in https://phytozome.jgi.doe.gov/pz/portal.html. To elucidate the evolutionary relationship across land plant evolutionary history, we focused on *A. thaliana, Selaginella moellendorffii, Physcomitrium patens* and *M. polymorpha*, as representative species for angiosperms, lycophytes, mosses and liverworts, respectively. Complete or partial coding protein sequences were aligned using the ClustalW parameters and were conducted in MEGA7. Phylogenetic trees were constructed with Neighbor-joining method, with a bootstrap test of 1000 replicates. The evolutionary distances were computed using the Poisson correction method.

### Vector construction

Expression constructs: a BJ36 plasmid [86] containing a tripleVENUS-NLS (trpVNS) fragment was modified by adding a ligation-independent cloning (LiC) adapter site [87]. A promoter fragment of Mp*FER* of 3.2 kb was amplified using primers specified below and cloned into the BJ36 vector via LiC cloning. *_pro_*Mp*FER* promoter fragment fused to trpVNS was then shuttled via *Not*I restriction sites into the HART01 [86] expression vector. The expression vector was also directly modified to contain the LiC sites, so promoter fragments can now be directly cloned in front of the trpVNS in the HART01 vector via LiC-cloning. This vector was called VHL (trpVNS-containing HART01 vector with LiC sites).

amiRNA constructs: For functional analyses three independent artificial microRNA (amiRNA) constructs were made (Fig. S2). The endogenous microRNA precursor Mp*miR160* was used as both backbone and template to design the constructs which generate single species small RNA molecules of 21 nt length, complementary to the target gene transcript [67], [88]. Three miR160 backbones containing independent amiR-Mp*FER*/amiR-Mp*FER*\* duplexes were synthesized by GenScript and fused to the *_pro_*Mp*EF1* promoter in the BJ36 shuttle vector and shuttled to the HART01 expression vector as described previously [88]. The folding structure and design procedures were explained in detail in [67], [88]. The sequences for the amiRNAs were designed to retain the exact physical properties of the endogenous Mp*miR160* template. The folding patterns were analyzed in the Mfold software [89] and the sequences synthesized (Fig. S2).

CRISPR/Cas gRNA: Selection of target sites was done using CasFinder [90]. Complementary oligos were designed and annealed for ligation into gRNA, in the pMpGE_En03 vector, previously digested with BsaI restriction enzyme. Resulting gRNAs were incorporated into binary vector pMpGE011 through gateway recombination.

Over-expression of Mp*FER* and At*FER*: The full-length Mp*FER* sequence was amplified from genomic *M. polymorpha* DNA with attB sites and Gateway-cloned via pDONR207. For *A. thaliana* expression, Mp*FER* was introduced into the expression vectors pMDC201, which contains a *2X35S* promoter fragment and an in-frame C-terminal mGFP6 [91]. For *M. polymorpha* expression, Mp*FER* was introduced intro the binary vector pMpGWB308, which contains a MpEF1 promoter and an in-frame C-terminal citrine [92]. Similarly, for expression of At*FER* in *M. polymorpha*, coding sequence of AtFER was introduced into pMpGWB308 binary vector, for the generation of *_pro_*Mp*EF1:*At*FER-Cit* construct.

### Quantitative real-time PCR (qRT-PCR) and droplet digital PCR (ddPCR)

RNA extraction from *M. polymorpha*: RNA extraction from thallus tissue (ca. 100mg including apical notch) was done using the Rneasy plant mini kit (#74904, Qiagen). To remove contaminating DNA, the TURBO DNAfree^TM^ Kit (AM1907 Ambion) was used according to manufacturer’s recommendations. To extract RNA from the gametophores (2 antheridiophores > 5mm diameter were pooled, respectively, for each replicate), the Direct(-zol) RNA MiniPrep (#R2050, ZymoResearch) was used with TRIzol Reagent (#15596026, Ambion) according to the manufacturer’s protocol, including on-column DNase digestion and subsequent TURBO DNA-free_TM_ Kit (AM1907 Ambion) treatment. RNA samples were quantified using the Qubit^®^ RNA HS Assay Kit (#Q32852, Life Technologies), a Nanodrop ND-1000 Spectrometer or an Agilent 2100 BioAnalyzer. cDNA synthesis: cDNA was synthesized from 0.5 *μ*g or 1.0 *μ*g of total RNA. Reverse transcription was performed in 25 *μ*l with 20*μ*g/ml Oligo(dT) 12-18 primers and 200 units of SuperScript^®^ II Reverse Transcriptase (#18064-014, Invitrogen) according to the manufacturer’s protocol. The resulting cDNA was diluted 1:9 by adding nuclease-free water. To control for genomic DNA contamination, control replicates of all samples were incubated without SuperScript^®^ II reverse transcriptase.

Primer tests and quantitative RT-PCR: Primer efficiency and concentration tests were carried out for Mp*FER* and suitable reference genes as described previously (Rövekamp et al., 2016, Althoff et al., 2014). Amplification experiments were carried out using a 7500 Applied Biosystem Fast Real-Time PCR System. Reactions were performed in 20 *μ*l volumes containing 10 *μ*l 2X SYBR-green mastermix (SsoAdvanced^TM^ Universal SYBRGreen Supermix), 200 nM (Mp*EF1* and Mp*FER*) or 250 nM (Mp*ACTIN*) forward and reverse primers, and 1 *μ*l diluted cDNA. Where possible, three technical and biological replicates were performed for each reaction. The primers used for the qRT-PCR are summarized in Table S1.

For ddPCR analysis, individual PCR reactions were performed in a total volume of 25 *μ*l, using 1X ddPCR EvaGreen Supermix, with droplets generated according to manufacturer’s recommendations. Reading of the PCR-amplified droplets was carried out by the QX200 Droplet Reader (Bio-Rad) and analysed by the QuantaSoft TM Software (v1.4, Bio-Rad). The counts from ddPCR were normalised through a log2 transformation. Afterwards, relative expression was calculated against the geometric mean of the counts for all three reference genes (Mp*ACT1*, Mp*ACT7*, and Mp*APT3*) [93]: (log2(gene tested + 1) −log2(geom. mean of all references + 1)). The primers used for the ddPCR are summarized in Table S1.

### 3’ RACE-PCR

RNA from wild-type and Mp*fer-1* mutant plants were extracted as previously described. A poly(G)-tail was added at the 3’ end of the RNA according to a Poly(A) Polymerase protocol (Thermo Scientific), using GTP instead of ATP. Reverse transcription was performed using anchored Oligo(dC) instead of Oligo(dT). Nested PCR on the cDNA was performed using two forward primers that matched the 3’ UTR sequence of Mp*FER* (3’RACE-Fwd1 and 3’RACE-Fwd2) and Oligo(dC). PCR fragments were purified and sequenced.

### Transformation of M. polymorpha and A. thaliana

*Agrobacterium*-mediated transformation using regenerating thalli of *M. polymorpha* was done by co-cultivation 15-day old plant fragments with transformed *Agrobacterium tumefaciens* (GV3101) cells [94], [95]. After three days of incubation, positive transformants were selected on Gamborg’s B5 plates supplemented with 10 *μ*g/ml Hygromicin B (Invitrogen), 0.5 μM Chlorosulfuron or Kanamycin, and 100 μg/ml of Cefotaxime sodium. Isogenic lines were obtained by using plants derived from gemmae of the T1 generation (G1 generation). G1 or subsequent gemmae generations were used for all experiments, as they are derived from single cells [96], [97]. Transformation of *A. thaliana* via *A. tumefaciens* (GV3101) was performed by floral dipping according to [98]. Primers used for amplification of promoter fragments of Mp*FER* are listed in Table S2.

### Microscopy

Plants were observed in a Lumar.V12 (Zeiss) or a Leica MZFLIII dissection microscope and photographed with an AxioCam HRc (Zeiss) or Leica DFC 420C camera. Clearings were observed using a Leica DMR microscope and photographed with an Axiocam 105 color camera. Fluorescence reporter expression were analyzed using a Leica SP5 confocal microscope. Fiji [99] and GIMP (version 2.8.10) software were used for adjustments of brightness, contrast, channel selection, and image size.

Tissue clearing for bright field microscopy: *M. polymorpha* tissues were fixed in Carnoy’s solution and incubated at 4°C overnight, followed by rehydration in an ethanol (EtOH) series of 85%, 70%, 50%, and 30%. Samples were incubated for 1 hour at 4°C for each step and then incubated in chloral hydrate solution at 4°C overnight. Samples were mounted in chloral hydrate.

Calcofluor-white staining for cell size measurements: Pieces of fresh thallus tissue located in fully differentiated zones [100] of plants grown on plates were dissected out and put into water. The pieces were transferred to PBS pH 6.1 containing 100 *μ*g/ml Calcofluor-white and vacuum-infiltrated for 1 hour before being mounted on slides. The area of the epidermal cells was measured using Imaris 8.3.1. software. Cells of air pores and the two adjacent cell layers were excluded from the analysis, as well as the spiralling cells surrounding newly developing air pores.

For cell-death staining, seven-day old gemmalings were stained with lactophenol-trypanblue (10 mL of lactic acid, 10 mL of glycerol, 10 g of phenol, 10 mg of trypan blue, dissolved in 10 mL of distilled water) and boiled for approximately 1 min in the stain solution and then decolorized in chloral hydrate (2.5 g of chloral hydrate dissolved in 1 mL of distilled water) for at least 30 min. They were mounted in chloral hydrate and imaged in dissecting microscope.

For scanning electron microscopy (SEM), 20-day old plants of WT and *_pro_*Mp*EF1:*Mp*FER-Cit* lines *#5* and #9 were harvested and fixed in using Glutaraldehyde (0.25% in 20mM HEPES) over night at 4 °C. After fixation, dehydratation with Acetone series (from 50% to 100% acetone “ultradry”) was done. The samples were then critical point dried, without the following sputter coated step. Microscopic imaging was done on a JSM-6010 (Jeol, Freising, Germany) at the Center for Microscopy and Image Analysis at the University of Zurich.

To evaluate the stiffness of the gemmaling tissue, we used cellular force microscopy (CFM; [101], [102]). A microelectromechanical systems (MEMS)-based FT-S100 force sensor (FemtoTools AG, Buchs, Switzerland) with a glued-on tungsten probe (T-4-22; Picoprobe by GGB Industries INC, Naples, FL, USA) was used to indent the gemmaling tissue. The sensor has a force range of ±100 μN and a resolution of 15 nN measured at 500 Hz. The sensor was attached with a custom-made acrylic arm to a xyz positioner (SLC-2460-M; SmarAct), which was fixed on a xyz piezo stage (P-563.3CD PIMars; Physik Instrumente (PI) GmbH & Co., Karlsruhe, Germany). The xyz positioner was used for the rough positioning of the sensor on a part of a gemma that was as much as possible in contact with the supporting glass slide and away from the meristematic zones. Nine evenly spaced measurements were then taken in a 3-by-3 grid, again using the xyz positioner to place the sensor, while the indentation at each position was driven by the piezo stage. This procedure was repeated a second time on each gemmaling, whenever possible on the opposite lobe.

The stiffness analysis was achieved with Matlab (Mathworks, Natick, MA, USA) based on the slope fitted to the force-displacement curves, which were acquired by CFM. In a first round of data curation, each force-indentation curve was visually inspected, and curves that didn’t have a clear contact point or showed a clear displacement of the gemmaling, indicated by an irregular slope, were discarded. The remaining data exhibited a bimodal distribution with a high density at very soft values around 1 µN, indicating the presence of an artefact due to gemmalings that were not in contact with the slide and, therefore, were displaced rather than indented. To identify such failed measurements, we pooled all data and fitted a Gaussian mixture model with two components. We then removed all measurements that were closer than 3 standard deviations to the mean of the distribution with the smallest mean (mean = 1.16 µN and sd = 0.56 µN). Hence, all measurements with a value below 2.9 µN were removed.

Boxplots and statistical analysis were performed in Matlab. The statistical tests used to assess the significance of differences between groups are indicated in the figure legends.

### Western blot analysis

*M. polymorpha* explants were ground in liquid nitrogen and resuspended in 300 μL of Laemmli buffer [103]. After centrifugation at 6,000 rpm for 5 min at 4°C, the supernatant was recovered. Protein accumulation was confirmed by Western blotting, using the anti-GFP antibody (1:3000, Torrey Pines Biolabs). As secondary antibodies, the Agrisera S09602 goat anti-rabbit antibody was used with the ECL-chemistry of FUSION FX -Western Blot & Chemi Imaging (Vilber Lourmat).

### Growth rate experiments

Gemmae of wild-type, Mp*fer-1*, and independent transformation lines of amiR-Mp*FER* lines were grown on plates and the thallus area was measured over time. 10 or 30 gemmae of four independent transformation lines were grown on half-strength Gamborg’s B5 and scanned on an Epson 2450 Photo scanner at 600dpi. Plant area was measured using color threshold settings and particle analysis in the Fiji software [99].

### Expression map

Gene expression of *Cr*RLK1L signaling pathway components was calculated from publicly available RNA-seq data of 39 samples of *M. polymorpha* Tak-1 and Tak.2. Samples were grouped by tissue and analyzed under the same pipeline: Nextera paired-end adapters were trimmed from sequencing reads using the bbduk tool embedded in the BBMap software package [104]. Read ends with quality below 20 were also trimmed and the minimum read length was set to 25. The rest of parameters of bbduk were set as default. Reads were mapped to *M. polymorpha* reference genome (v3.1) [48] using the Tophat software (v2.1.1) [105], designed for RNA-sequencing read mapping. Tophat contains Bowtie (v2.3.2.0) [106] as the aligner software. All parameters were set to default values except for the RAM memory and number of used Cores. Mapped reads were sorted using samtools (v1.3.1) [107] and counted with the software HTSeq-count” (v.0.9.1) [108]. Gene expression values were calculated using the package DESeq2 (Release 3.6) [109] for R software (v3.4.4) [110]. Another R package, ggplot2 (release 2.2.1) [111] was used for producing the figures. A summary of the samples that were used in this study is provided in Table S3.

### Chlorophyll efflux rate

To measure the chlorophyll efflux rate, 10-day-old gemmalings were used [112]. Sixteen replicates of MpFER-overexpressor, Mp*fer*-2 mutant, and wild-type lines were immersed in an 80% EtOH solution in 5-mL tubes. The tubes were agitated gently on a shaker platform. Aliquots of 2 µL were removed every min, during the first 10 minutes. The amount of chlorophyll extracted by the EtOH solution was quantified with a spectrophotometer (Nanodrop 8000, Thermo Scientific) and calculated from UV light absorption at 647 and 664 nm. The micromolar concentration of total chlorophyll per gram of fresh weight of tissue per ml of EtOH was calculated using the equation Total micromoles chlorophyll = 7.93(A_664_) + 19.53(A_647_).

### Three-dimensional (3D) protein structure modeling

Comparative modeling of protein 3D structures was performed using the SWISS-MODEL online-tool [64]. The software was fed with the three protein sequences of MpFER, MpRALF1, and MpLRE1 as input to model, and one PDB file as a 3D template, which contains the published crystal structure of the protein complex formed by AtFER, AtRALF23, and AtLLG1 [36]. Quality control of the model was assessed using the same tool and the results are summarized in Table S4. Spatial comparisons of the *M. polymorph*a predicted structures with the crystal structures of *A. thaliana* were performed in PyMol (Table S4) [113].

## QUANTIFICATION AND STATISTICAL ANALYSIS

Analyses were performed in InfoStat (http://www.infostat.com.ar) or Matlab. Tests are indicated in the corresponding figure legend.

## Acknowledgments

We thank Liam Dolan for kindly providing the Mp*fer-1*/Mp*the* mutant, Valeria Gagliardini for help with qRT-PCR and ddPCR, Anja Grossniklaus for help in measuring epidermal cell area, Célia Baroux and Ethel Mendocillo-Sato for instructions on confocal microscopy and the use of Imaris Software, Tiago Meier for helping with sample preparation for SEM microscopy, Isabel Monte and Cyril Zipfel for comments on the manuscript, and Christof Eichenberger, Frédérique Pasquer, Arturo Bolaños, Daniela Guthörl, Daniel Prata, and Peter Kopf for general lab support.

## Author contributions

MAM: Conceptualization, Formal analysis, Investigation, Methodology, Validation, Visualization, Writing – original draft, Writing – review & editing

MR: Conceptualization, Formal analysis, Investigation, Methodology, Validation, Visualization, Funding acquisition, Writing – original draft, Writing – review & editing

AG-F: Data curation, Formal analysis, Software, Validation, Visualization, Writing – original draft, Writing – review & editing

DL: Investigation, Validation

DM: Formal analysis, Investigation, Validation, Writing – review & editing

PG: Formal analysis, Investigation, Validation, Writing – review & editing

HV: Formal analysis, Investigation, Validation, Writing – review & editing

JLB: Funding acquisition, Resources, Supervision, Validation, Writing – review & editing

UG: Conceptualization, Formal analysis, Funding acquisition, Project administration, Supervision, Validation, Writing – original draft, Writing – review & editing

