## Supplementary information for "The single *Marchantia polymorpha FERONIA* homolog reveals an ancestral role in regulating cellular expansion and integrity"

**A**

| | MpFER | | |
|---|---|---|---|
| **% Identity** | Malectin | ECD | Full protein |
| AtFER | 33.5 | 31.0 | 43.3 |
| AtHERK2 | 31.7 | 31.8 | 45.7 |
| AtANX2 | 30.6 | 27.2 | 40.3 |
| AtANX1 | 29.8 | 26.5 | 39.8 |
| AtTHE1 | 29.2 | 28.4 | 46.0 |

**B**

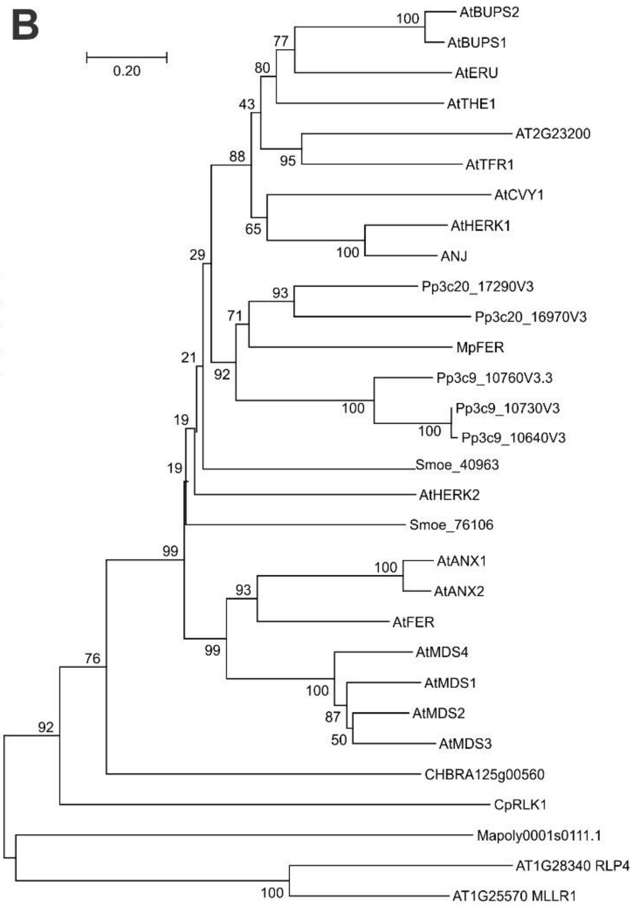

**C**

| **% Identity** | ECD CpRLK1 | CpRLK1 |
|---|---|---|
| MpFER | 25.7 | 29.2 |
| Mp1g17720 | 35.3 | 45.7 |
| AtFER | 28.0 | 41.5 |

**D**

| **RMSD** | AtFER |
|---|---|
| MpFER | 0.139 |
| CpRLK1 | 2.347 |
| CHBRA125g00560 | 0.352 |

**E**

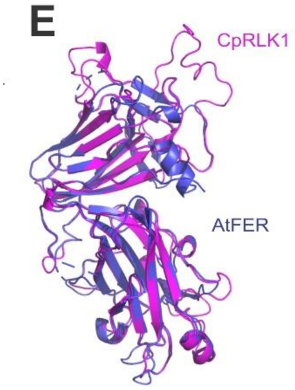

**F**

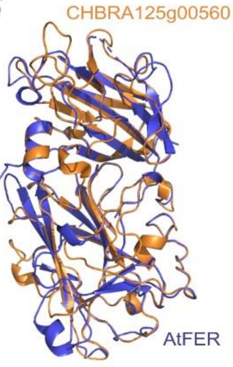

**Figure S1. The *Cr*RLK1L Family Appeared together with Land Plants. Related to Fig. 1.**

(A) Percentage of identity of the malectin-like domain, the complete ECD, and the full protein of MpFER with AtFER, AtHERK2, AtANX2, AtANX1, and AtTHE1.

(B) A rooted neighbour-joining tree of the amino acid sequence of the predicted malectin-like domains was generated using ClustalW. *Cr*RLK1L members from *Marchantia polymorpha* (Mp), *Physcomitrium patens* (Pp), *Selaginella moellendorffii* (Smoe), and *Arabidopsis thaliana* (At) were used. Algal CpRLK1 and CHBRA125g00560, as well as Mp1g17720 (Mapoly001s0111) were also included. The numbers indicate the bootstrap values (%) from 1000 replications. The given scale represents a substitution frequency of 0.1 amino acids per site.

(C) Percentage of identity of the ECD and full protein of CpRLK1 with MpFER, AtFER, and Mp1g17720 (Mapoly001s0111).

(D) RMSD values for prediction of MpFER, CpRLK1, and CHBRA125g00560 3D structures of the ECD based on the AtFER ECD.

(E) Structural superposition of the ECD of AtFER (blue) and CpRLK1 (purple).

(F) Structural superposition of the ECD of AtFER (blue) and CHBRA125g00560 (orange) from *Chara brunii*.

**A**

Mp*miR160* Mp1g26670
GCACCTCCTCTCTCCGACTGCAGCCCGTTTCGAGATCCGAGGACTTGCTCGACGCGACTAATTGGGGAGGCCAGACTG
CACTTGCCTGGCTCCCTGTATGCCAACTGAGGAGCTCCTCAGAGACCTTGACAGGCTCCGTAGCTGGCATTCAGGGGG
CCATGCAGGAGGAAGTCGCTACCTCCCGCAAGGTGCGACTAGCTTTCTGTCTTGGGTGCACACCTCACTGATGTTTGA
TAGATTTACTTA

amiR-Mp*FER*1
**TGCAAGCTT**GCACCTCCTCTCTCCGACTGCAGCCCGTTTCGAGATCCGAGGACTTGCTCGACGCGACTAATTGGGGAG
GCCAGACTGCACTTGACGTAGGCAAACTTGTCGAACTGAGGAGCTCCTCAGAGACCTTGACAGGCTCCGTAGCTCGAC
ATGTTTGTCTACCTCAGGAGGAAGTCGCTACCTCCCGCAAGGTGCGACTAGCTTTCTGTCTTGGGTGCACACCTCACT
GATGTTTGATAGATTTACTTA**GGATCCATA**

amiR-Mp*FER*2
**TGCAAGCTT**GCACCTCCTCTCTCCGACTGCAGCCCGTTTCGAGATCCGAGGACTTGCTCGACGCGACTAATTGGGGAG
GCCAGACTGCACTTCGATGTCGAAAATCCAGGTGACTGAGGAGCTCCTCAGAGACCTTGACAGGCTCCGTAGCCACCT
GCATTTTTGACAACGAGGAGGAAGTCGCTACCTCCCGCAAGGTGCGACTAGCTTTCTGTCTTGGGTGCACACCTCACT
GATGTTTGATAGATTTACTTA**GGATCCATA**

amiR-Mp*FER*3
**TGCAAGCTT**GCACCTCCTCTCTCCGACTGCAGCCCGTTTCGAGATCCGAGGACTTGCTCGACGCGACTAATTGGGGAG
GCCAGACTGCACTTCCGAACTTCCTCAGAGACTCACTGAGGAGCTCCTCAGAGACCTTGACAGGCTCCGTAGCGAGTC
TGTGAGGGAGTTGGGAGGAGGAAGTCGCTACCTCCCGCAAGGTGCGACTAGCTTTCTGTCTTGGGTGCACACCTCACT
GATGTTTGATAGATTTACTTA**GGATCCATA**

**B**

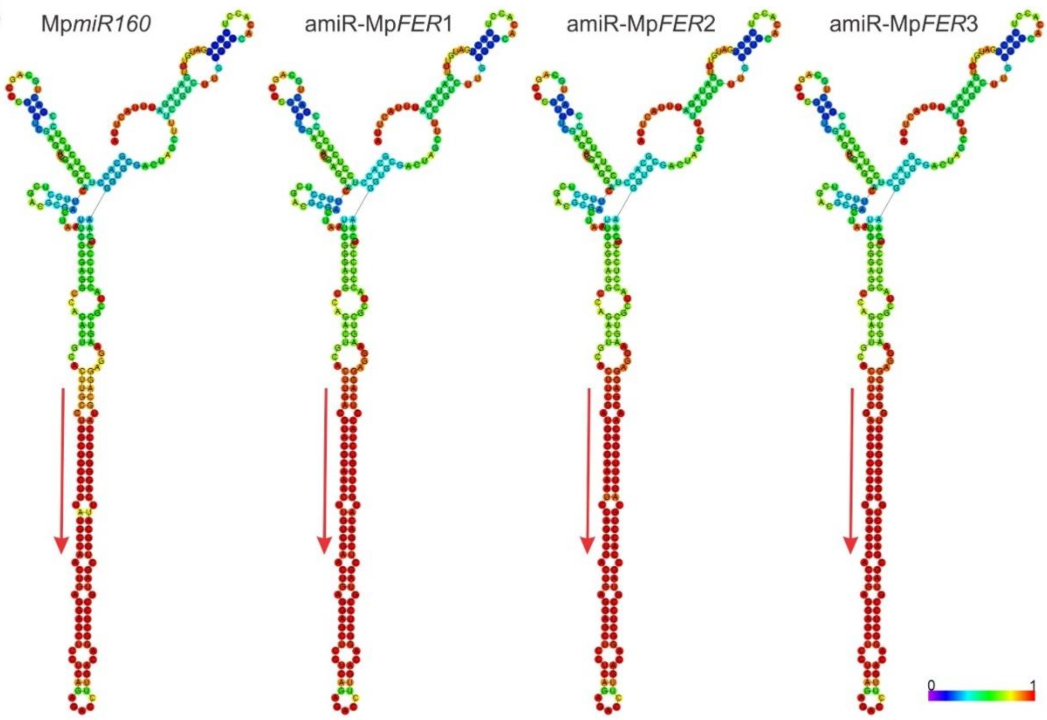

**Figure S2. Design of amiR-Mp*FER* Precursors. Related to Fig. 3.**

(A) Mp*miR160* (Mapoly0002s0211, Mp1g26670) and amiR-Mp*FER*1, amiR-Mp*FER*2, and amiR-Mp*FER* sequences. miRNA sequences are in red and miRNA* in blue. Cloning sequences from amiR-Mp*FER* constructs are in bold.

(B) Drawing of the minimum free energy structure of Mp*miR160* and amiR-Mp*FER*3 constructs predicted by the RNAfold web server (http://rna.tbi.univie.ac.at//cgibin/RNAWebSuite/RNAfold.cgi). Red arrows indicate location and orientation of the mature miRNA in the precursor. The structures are coloured by base-pairing probabilities; for unpaired regions the colour denotes the probability of being unpaired.

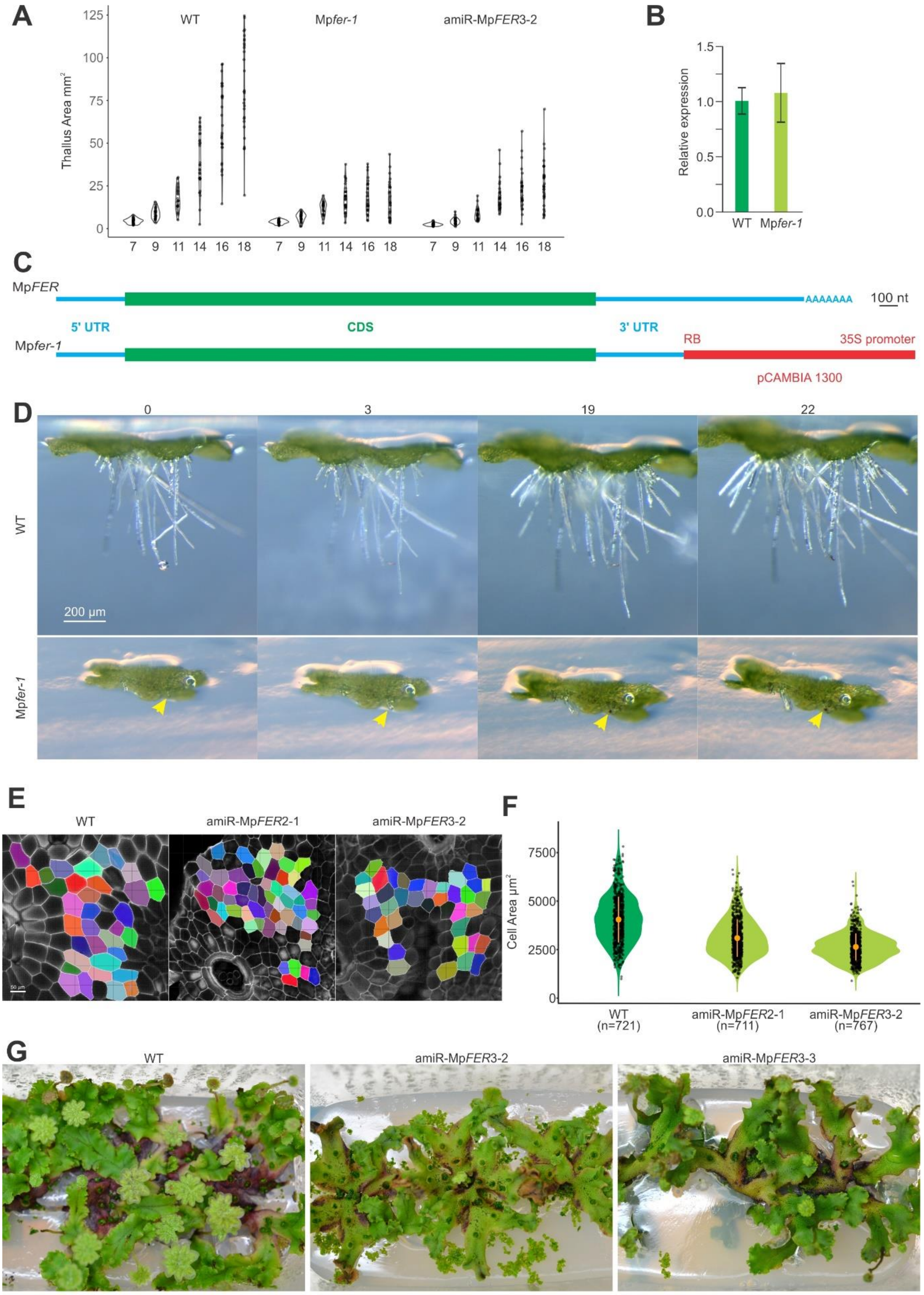

**Figure S3. Reduction in Thallus and Cell Area in Plants with Reduced Mp*FER* Levels. Related to Figs. 3 and 5.**

(A) Violin plot of thallus area of wild-type (WT), Mp*fer-1*, and amiR-Mp*FER*3-2 plants at different days after putting gemmae on plates. n = 30. Areas were estimated using ImageJ software.

(B) Relative expression level of Mp*FER* in 14 day-old WT and Mp*fer-1* gemmalings, as measured by qRT-PCR. Mp*EF1* was used as internal control. Shown are means ± standard error of the mean (SEM) of three biological replicates. Statistical analysis was performed by a one-way analysis of variance (ANOVA) follow by a post-hoc Duncan test, no significant difference was observed.

(C) Schematic of Mp*FER* transcripts in WT (upper) and Mp*fer-1* plants (lower).

(D) Pictures of growing rhizoids at different time points (in h). Yellow arrows indicate a growing rhizoid in Mp*fer-1* line that burst between 3 and 19 h.

(E) Representative images from cell surface areas measured in WT, amiR-Mp*FER*2-1, and amiR-Mp*FER*3-2 plants. Scale bar, 50 *µ*m.

(F) Violin plot of cell areas of WT, amiR-Mp*FER*2-1, *and* amiR-Mp*FER*3-2 plants. Difference is significant based on the nonparametric Kruskal-Wallis test and a linear regression model with a highly significant interaction ($p < 0.001$). Orange circles indicate the group mean and the corresponding vertical bars the standard deviation for each group.

(G) Induction of antheridiophores in WT and two independent amiR-Mp*FER*3 lines. Three plants were grown in each sterile plastic box under far-red light induction.

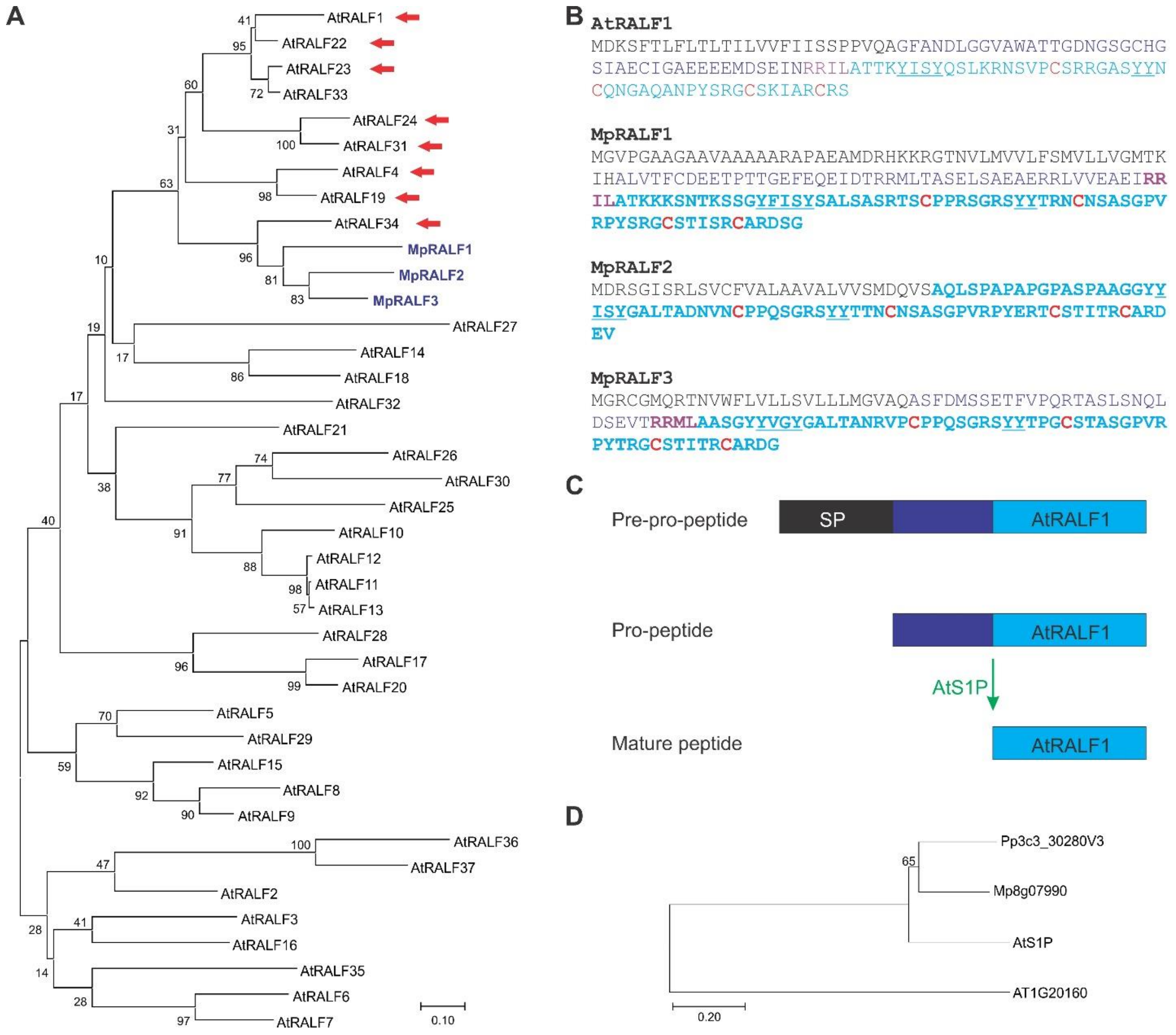

**Figure S4. MpRALF Peptides Belong to the AtRALF1-clade of the RALF Family. Related to Fig. 6.**

(A) A rooted neighbour-joining tree of the amino acid sequence of the predicted mature RALF peptides was generated using ClustalW. RALF members from *M. polymorpha* and *A. thaliana* were used. Red arrows indicate RALFs that are known ligands of *Cr*RLK1Ls. The numbers indicate the bootstrap values (%) from 1000 replications. The given scale represents a substitution frequency of 0.1 amino acids per site.

(B) Amino acid sequence comparison of AtRALF1 and MpRALF1-3. Predicted signal peptides are in black, predicted mature peptides in light blue, conserved Cys in red, and predicted S1P recognition sites in violet.

(C) Processing pathway AtRALF1 by the S1P protease.

(D) A rooted neighbour-joining tree of the amino acid sequence of the S1P orthologs was generated using ClustalW. S1P members from *M. polymorpha*, *P. patens*, and *A. thaliana* were used. The numbers indicate the bootstrap values (%) from 1000 replications. The given scale represents a substitution frequency of 0.1 amino acids per site.

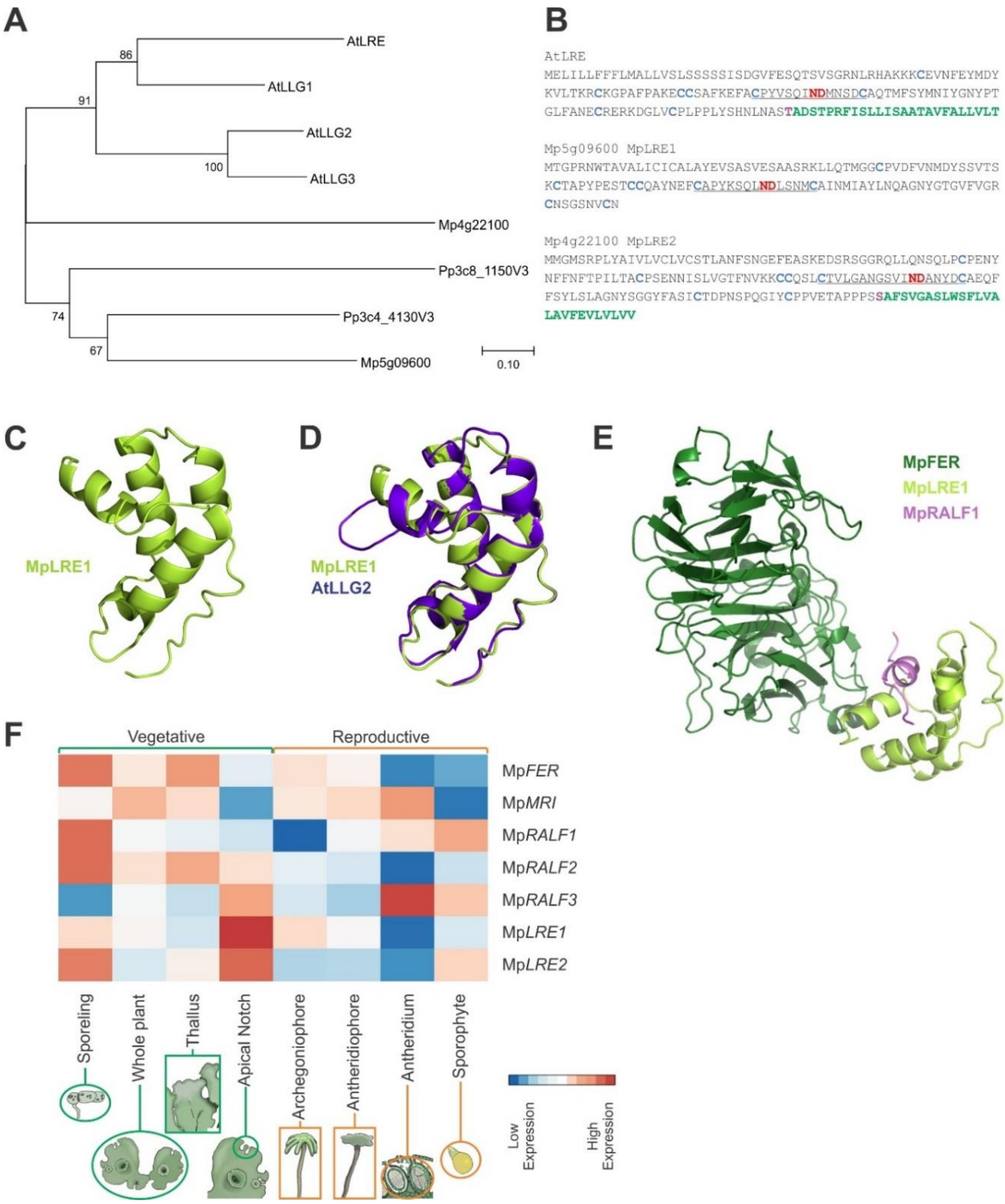

**Figure S5. Two LORELEI-like Proteins Are Encoded in the *M. polymorpha* Genome. Related to Fig. 6.**

(A) A rooted neighbor-joining tree of the amino acid sequence of LRE orthologs was generated using ClustalW. LRE members from *M. polymorpha*, *P. patens*, and *A.*

*thaliana* were used. The numbers indicate the bootstrap values (%) from 1000 replications. The given scale represents a substitution frequency of 0.1 amino acids per site.

(B) Amino acid sequence of AtLRE, MpLRE1, and MpLRE2. Conserved Cys are in light blue, the ND motif in red, and the GPI-anchoring site in green.

(C) Cartoon representation of the predicted 3-dimensional structure of MpLRE1, showing predicted alpha-helices.

(D) Structural superposition of AtLGG2 (blue) and MpLRE1 (green).

(E) Cartoon representation of the predicted 3-dimensional structure of the MpFER/MpLRE1/MpRALF1 complex, showing predicted alpha-helices and beta-sheets.

(F) Heatmap depicting relative gene expression based on RNAseq data (row-Z-score of vs normalized counts) of Mp*FER* and the *M. polymorpha* orthologs of At*MR*I, At*RALF1*, and At*LRE* across different tissues*.* Vegetative and reproductive tissues are grouped by green and orange, respectively. Averaged expression values are represented with colours of increasing red and blue intensity indicating upregulation and downregulation of gene expression, respectively.

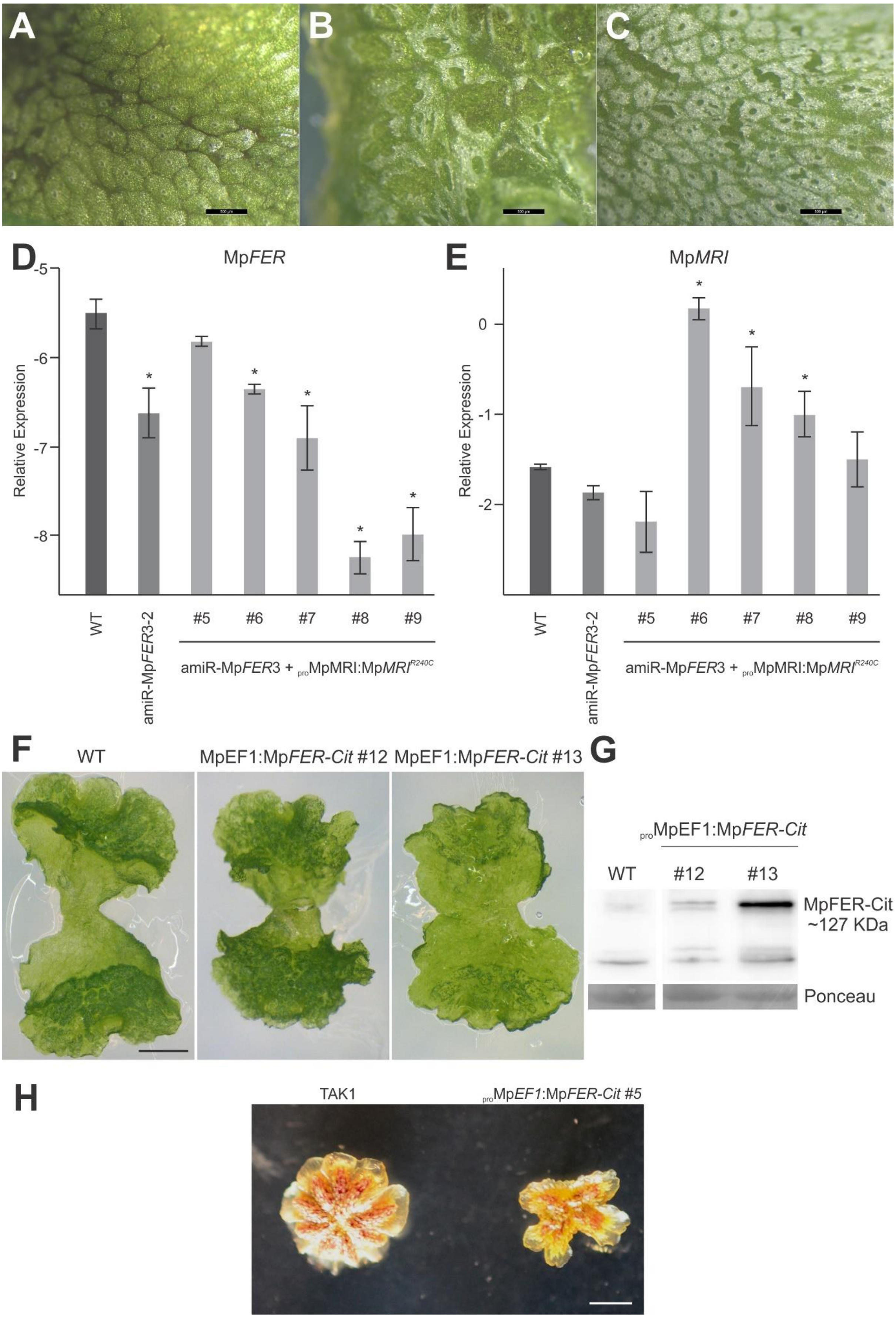

A
B
C
D
MpFER
Relative Expression
-5
-6
-7
-8
WT
amiR-MpFER3-2
#5
#6
#7
#8
#9
amiR-MpFER3 + proMpMRI:MpMRI R240C
E
MpMRI
Relative Expression
0
-1
-2
WT
amiR-MpFER3-2
#5
#6
#7
#8
#9
amiR-MpFER3 + proMpMRI:MpMRI R240C
F
WT
MpEF1:MpFER-Cit #12
MpEF1:MpFER-Cit #13
G
proMpEF1:MpFER-Cit
WT
#12
#13
MpFER-Cit
~127 KDa
Ponceau
H
TAK1
proMpEF1:MpFER-Cit #5

**Figure S6. Expression of Mp*MRI*$^{R240C}$ Suppresses the Bursted Rhizoid Phenotype of amiR-Mp*FER*3 Lines but Leads to Aberrant Epidermis Development. Related to Figs. 6,7.**

(A to C) Epidemical pictures of thalli from the wild-type (WT) (A), and amiR-Mp*FER*3 + $_{pro}$MpMRI:Mp*MRI*$^{R240C}$ lines #6 (B) and #8 (C), which both partially suppressed the bursting rhizoid phenotype (Fig. 6).

(D and E) Relative expression of Mp*FER* (D) and Mp*MRI* (E) against the geometric mean of the reference genes Mp*ACT1*, Mp*ACT7*, and Mp*APT3* in WT, amiR-Mp*FER*3-2, and 5 lines (#5 to #9) co-transformed with the amiR-Mp*FER*3-2 and $_{pro}$MpMRI:Mp*MRI*$^{R240C}$ constructs. Expression levels of three biological replicates were assessed by droplet digital PCR (ddPCR). The y-axis corresponds to the log2-ratio between the test and the geometric mean of the reference genes. Shown are means ± SEMs of three biological replicates. Statistical analysis was performed by one-way analysis of variance (ANOVA) followed by a post-hoc Duncan test (*P < 0.01).

(F) Representative pictures of 10-day old gemmalings of wild-type (WT) and two different lines overexpressing Mp*FER* ($_{pro}$Mp*EF1:*Mp*FER-Cit)*. Scale bar, 1 mm.

(G) Western blot analysis of $_{pro}$Mp*EF1:*Mp*FER-Cit* lines from Fig. S6A using an anti-GFP antibody. WT lines were used as negative controls. The Ponceau membrane staining of the most intense band at 55 kDa (presumably Rubisco) was used as a loading control.

(H) Representative images of the antheridial receptacle of WT and $_{pro}$Mp*EF1:*Mp*FER-Cit* plants. Scale bar, 2 mm.

| Name | Sequence |
|---|---|
| Mp*FER* | CGAGGAGCATTGCGAGATG |
| Mp*FER* | AGGTCGGTGCCGTAGAGATG |
| Mp*EF1α* | AGGTTGTCACCATGGGAAAGGAGA |
| Mp*EF1α* | TCACACGCTTGTCAATACCTCCCA |
| Mp*MRI* | TGGCAGCTCGTCTCCACTCT |
| Mp*MRI* | AGTCATGGCGTACTCGGGTG |
| Mp*ACT7* | AGGCATCTGGTATCCACGAG |
| Mp*ACT7* | ACATGGTCGTTCCTCCAGAC |
| Mp*ACT1* | GAGCGCGGTTACTCTTTCAC |
| Mp*ACT1* | GACCGTCAGGAAGCTCGTAG |
| Mp*APT3* | CGAAAGCCCAAGAAGCTACC |
| Mp*APT3* | GTACCCCCGGTTGCAATAAG |
| 3'RACE-Fwd1 | AACGGTGGTTGGATGGTCGATTAG |
| 3'RACE-Fwd2 | CGACATGGACGGCCTGGAGCTGGAG |
| Oligo(dC) | CCCCCCCCCCCCCCCCCVN |

**Table S1: Primer sequences used for qRT-PCR, ddPCR, and 3' RACE-PCR**

| Name | Target | Sequence |
|---|---|---|
| $_{pro}$MpFER F | Mp*FER* | TAGTTGGAATGGGTTCGAATGCTGTCGACCACTGACTTC |
| $_{pro}$MpFER R | Mp*FER* | TTATGGAGTTGGGTTCGAAGTAGTGTATCCTCCAGCCGCTTT |
| MpFER F | Mp*FER* | GGGGACAAGTTTGTACAAAAAAGCAGGCT-AGAGCCCAAGGAGGAAGGGCGACCA |
| MpFER R | Mp*FER* | GGGGACCACTTTGTACAAGAAAGCTGGGTT-CCTTCCTTGAGGGTTCACCAGCTG |

**Table S2: List of primers used for cloning**

| SRR_ID | SRX_ID | Organism | Tissue Group | Phase | Strain |
|---|---|---|---|---|---|
| **SRR896228** | SRX301558 | M.Polymorpha;WT | Thallus | Vegetative | Tak1 |
| **SRR896225** | SRX301555 | M.Polymorpha;WT | Archegoniophore | Sexual | Tak2BC4 |
| **SRR896226** | SRX301560 | M.Polymorpha;WT | Thallus | Vegetative | Tak1 |
| **SRR896229** | SRX301557 | M.Polymorpha;WT | Thallus | Vegetative | Tak1 |
| **SRR896224** | SRX301559 | M.Polymorpha;WT | Sporeling | Vegetative | Tak1xTak2BC4 |
| **SRR896230** | SRX301553 | M.Polymorpha;WT | Antheridiophore | Sexual | Tak1 |
| **SRR896227** | SRX301554 | M.Polymorpha;WT | Thallus | Vegetative | Tak1xTak2BC4 |
| **SRR896223** | SRX301556 | M.Polymorpha;WT | Sporophyte | Sexual | Tak1xTak2BC4 |
| **SRR971246** | SRX346276 | M.Polymorpha;WT | Archegoniophore | Sexual | Tak2 |
| **SRR971244** | SRX346274 | M.Polymorpha;WT | Thallus | Vegetative | Tak2 |
| **SRR971248** | SRX346277 | M.Polymorpha;WT | Antheridiophore | Sexual | Tak1 |
| **SRR971249** | SRX346278 | M.Polymorpha;WT | Archegoniophore | Sexual | Tak2 |
| **SRR971243** | SRX346272 | M.Polymorpha;WT | Thallus | Vegetative | Tak1 |
| **SRR971245** | SRX346275 | M.Polymorpha;WT | Antheridiophore | Sexual | Tak1 |
| **SRR1553299** | SRX682817 | M.Polymorpha;WT | Sporophyte | Sexual | Tak1xTak2 |
| **SRR1552617** | SRX682160 | M.Polymorpha;WT | Whole_Plant | Vegetative | Tak1xTak2 |
| **SRR1553294** | SRX682811 | M.Polymorpha;WT | Apical_Notch | Vegetative | Tak1xTak2 |
| **SRR1553297** | SRX682815 | M.Polymorpha;WT | Sporophyte | Sexual | Tak1xTak2 |
| **SRR1553296** | SRX682814 | M.Polymorpha;WT | Apical_Notch | Vegetative | Tak1xTak2 |
| **SRR1553276** | SRX682793 | M.Polymorpha;WT | Whole_Plant | Vegetative | Tak1xTak2 |
| **SRR1553295** | SRX682813 | M.Polymorpha;WT | Apical_Notch | Vegetative | Tak1xTak2 |
| **SRR1553298** | SRX682816 | M.Polymorpha;WT | Sporophyte | Sexual | Tak1xTak2 |
| **DRR050343** | DRX045349 | M.Polymorpha;WT | Whole_Plant | Vegetative | Tak1xTak2 |
| **DRR050346** | DRX045352 | M.Polymorpha;WT | Antheridiophore | Sexual | Tak1xTak2 |
| **DRR050347** | DRX045353 | M.Polymorpha;WT | Antheridiophore | Sexual | Tak1xTak2 |
| **DRR050348** | DRX045354 | M.Polymorpha;WT | Antheridiophore | Sexual | Tak1xTak2 |
| **DRR050353** | DRX045359 | M.Polymorpha;WT | Archegoniophore | Sexual | Tak1xTak2 |
| **DRR050344** | DRX045350 | M.Polymorpha;WT | Whole_Plant | Vegetative | Tak1xTak2 |
| **DRR050351** | DRX045357 | M.Polymorpha;WT | Archegoniophore | Sexual | Tak1xTak2 |
| **DRR050349** | DRX045355 | M.Polymorpha;WT | Antheridium | Sexual | Tak1xTak2 |
| **DRR050352** | DRX045358 | M.Polymorpha;WT | Archegoniophore | Sexual | Tak1xTak2 |
| **DRR050345** | DRX045351 | M.Polymorpha;WT | Whole_Plant | Vegetative | Tak1xTak2 |
| **DRR050350** | DRX045356 | M.Polymorpha;WT | Antheridium | Sexual | Tak1xTak2 |
| **DRR118950** | DRX111959 | M.Polymorpha;WT | Whole_Plant | Vegetative | Tak1 |
| **DRR118945** | DRX111954 | M.Polymorpha;WT | Whole_Plant | Vegetative | Tak2BC3 |
| **DRR118951** | DRX111960 | M.Polymorpha;WT | Whole_Plant | Vegetative | Tak1 |
| **DRR118943** | DRX111952 | M.Polymorpha;WT | Whole_Plant | Vegetative | Tak2BC3 |
| **DRR118944** | DRX111953 | M.Polymorpha;WT | Whole_Plant | Vegetative | Tak2BC3 |
| **DRR118949** | DRX111958 | M.Polymorpha;WT | Whole_Plant | Vegetative | Tak1 |

**Table S3. List of publicly available RNA-seq samples downloaded from SRA and used in this study.** Sample list of publicly available RNA-seq samples that were used in this study. The table also includes the tissue classification in which every sample was grouped with, their corresponding phase (vegetative vs sexual), and the genotype they correspond two. BC = Back Crossed to TAK1 genotype followed by the number of back crosses.

| Template Gene | Target Gene | Template PDB | GMQE | QMEAN DisCo | RMSD |
|---|---|---|---|---|---|
| **AtFER** | MpFER | 6a5b.1.A | 0.66 | 0.71 ± 0.05 | 0.139 |
| **AtANX1** | MpFER | 6fig.1.A | 0.64 | 0.68 ± 0.05 | 0.144 |
| **AtANX2** | MpFER | 6fih.1.A | 0.58 | 0.62 ± 0.05 | 0.104 |
| **AtLLG1** | MpLRE1 | 6a5d.2.A | 0.52 | 0.70 ± 0.10 | 0.11 |
| **AtLLG2** | MpLRE1 | 6a5e.1.B | 0.45 | 0.63 ± 0.11 | 0.14 |
| **AtLLG1** | MpLRE2 | 6a5d.2.A | 0.44 | 0.67 ± 0.05 | 0.15 |
| **AtLLG2** | MpLRE2 | 6a5e.1.B | 0.34 | 0.57 ± 0.09 | 0.18 |
| **AtFER** | CpRLK1 | 6a5b.1.A | 0.45 | 0.51 ± 0.05 | 2.347 |
| **AtANX1** | CpRLK1 | 6fig.1.A | 0.48 | 0.52 ± 0.05 | 3.038 |
| **AtANX2** | CpRLK1 | 6fih.1.A | 0.49 | 0.52 ± 0.05 | 1.721 |
| **AtFER** | CHBRA125g00560 | 6a5b.1.A | 0.58 | 0.61 ± 0.05 | 0.352 |
| **AtANX1** | CHBRA125g00560 | 6fig.1.A | 0.59 | 0.59 ± 0.05 | 0.41 |
| **AtANX2** | CHBRA125g00560 | 6fih.1.A | 0.59 | 0.59 ± 0.05 | 0.481 |

**Table S4. Quality estimation values generated during the modelling processes.** For each target protein, several templates were considered for model comparison and validation. General Model Quality Estimate (GMQE) score and the QMEAN DIsCo score are generated by SWISS-MODEL web server [60]: higher values indicate better accuracy in model prediction. RMSD stands for root-mean-square deviation and measures the average distance between the atoms of paired superimposed proteins, thus, the lower the values, the more similar is the modelled protein to the template.
